# Induced pluripotent stem cell-derived extracellular vesicles promote wound repair in a diabetic mouse model via an anti-inflammatory immunomodulatory mechanism

**DOI:** 10.1101/2023.03.19.533334

**Authors:** Daniel Levy, Sanaz Nourmohammadi Abadchi, Niloufar Shababi, Mohsen Rouhani Ravari, Nicholas H. Pirolli, Cade Bergeron, Angel Obiorah, Farzad Mokhtari-Esbuie, Shayan Gheshlaghi, John M. Abraham, Ian M. Smith, Emily Powsner, Talia Solomon, John W. Harmon, Steven M. Jay

## Abstract

Extracellular vesicles (EVs) derived from mesenchymal stem/stromal cells (MSCs) have recently been widely explored in clinical trials for treatment of diseases with complex pathophysiology. However, production of MSC EVs is currently hampered by donor-specific characteristics and limited *ex vivo* expansion capabilities before decreased potency, thus restricting their potential as a scalable and reproducible therapeutic. Induced pluripotent stem cells (iPSCs) represent a self-renewing source for obtaining differentiated iPSC-derived MSCs (iMSCs), circumventing both scalability and donor variability concerns for therapeutic EV production. Thus, we initially sought to evaluate the therapeutic potential of iMSC EVs. Interestingly, while utilizing undifferentiated iPSC EVs as a control, we found that their vascularization bioactivity was similar and their anti-inflammatory bioactivity was superior to donor-matched iMSC EVs in cell-based assays. To supplement this initial *in vitro* bioactivity screen, we employed a diabetic wound healing mouse model where both the pro-vascularization and anti-inflammatory activity of these EVs would be beneficial. In this *in vivo* model, iPSC EVs more effectively mediated inflammation resolution within the wound bed. Combined with the lack of additional differentiation steps required for iMSC generation, these results support the use of undifferentiated iPSCs as a source for therapeutic EV production with respect to both scalability and efficacy.

## 1. Introduction

While cell-based therapeutics featuring multipotent or progenitor cells have received significant interest in regenerative medicine and tissue repair applications ^1, 2^, research has demonstrated that secreted factors such as cytokines, chemokines, and, especially, extracellular vesicles (EVs), play a substantial role in their therapeutic effects ^3^. EVs are cell-secreted, naturally-occurring nanoscale particles that function in intercellular communication via transfer of nucleic acids, lipids, and proteins to recipient cells ^4^. The biomolecular composition of EV cargos is determined by their parental cells, and EVs derived from cell sources with therapeutic potential such as multipotent or progenitor cells possess many of the same regenerative properties ^3, 5^. Additionally, EVs are an attractive alternative to cell-based therapies due to a preferred safety profile as a result of their inability to replicate as well as their more predictable pharmacokinetic properties ^6, 7^. However, scalable production of both cell- and EV-based therapies is currently a key limitation to their clinical translation ^8, 9^.

Specifically, most cells used to produce therapeutic EVs have limited expansion capabilities ^10, 11^. This includes mesenchymal stem/stromal cells (MSCs), which are among the most widely utilized cell sources for therapeutic EV production due to their multifactorial regenerative properties ^10, 12^. Additionally, it has been demonstrated that increased *ex vivo* expansion of MSCs can affect their phenotype and therefore therapeutic efficacy; previously, it has also been observed that this decrease in efficacy translates to their secreted EVs ^10, 13^. Currently, sourcing adult MSCs from various donors is a feasible workaround to the issues with limited expansion ^14^. However, donor variance in age, sex, and other genetic differences creates significant variability in the therapeutic potency of MSCs and their secreted EVs ^15, 16^.

Towards addressing this limitation, researchers have attempted to develop a scalable source for therapeutic EVs by immortalizing MSCs ^17-19^. However, there are safety concerns associated with this strategy, as immortalization can make MSCs genetically similar to cancer cells ^18, 20^. Another approach is to utilize MSCs differentiated from self-renewing induced pluripotent stem cells (iPSC-MSCs, or iMSCs) ^21^, which can continually be produced from the same donor line, thus alleviating donor variability and scalability concerns, but at the cost of increased production time ^22^. Interestingly, researchers have begun to demonstrate the therapeutic utility of EVs from undifferentiated iPSCs, which require fewer processing steps to generate than iMSCs and thus are more favorable with respect to cost and reproducibility ^22, 23^. For example, *Adamiak* et al. were able to utilize iPSC EVs to improve cardiac function in mice post-myocardial infarction, while *Povero* et al. demonstrated that iPSC EVs partially reverse murine liver fibrosis ^24, 25^. While these initial works are promising and iPSC EV research is currently growing rapidly, studies have yet to benchmark the therapeutic efficacy of iPSC EVs against more established primary MSC or iMSC EVs.

In this work, we demonstrate that iPSC EVs possess similar pro-angiogenic bioactivity to donor-matched iMSC EVs *in vitro*. Additionally, for the first time, we demonstrate that iPSC EVs possess anti-inflammatory properties comparable or superior to iMSC EVs. Further, in a diabetic murine wound healing model, we show that when compared to donor-matched iMSC EVs, iPSC EVs have superior therapeutic properties, functioning via modulation of the immune infiltrate. These results demonstrate that iPSC EVs may be a feasible therapeutic modality in tissue repair applications that require simultaneous modulation of complex, multifunctional regenerative pathways.

## 2. Methods

### 2.1 Cell culture

Human iPSCs (ACS-1026; American Type Culture Collection, Manassas, VA, USA) were cultured in mTESR Plus (100-0276; STEMCELL Technologies, Cambridge, MA, USA) complete medium on hESC-qualified Matrigel basement matrix (35277; Corning; Corning, NY, USA) in either cell culture treated 6-well plates or T-75 tissue culture flasks; iPSCs arrived from the manufacturer at passage 22 and were not used for EV production for functional assays after more than 35 total passages. iPSCs were passaged before colonies began to touch and differentiate. Large particle-depleted mTESR Plus was generated by centrifugation of the complete medium at 100,000 x g for 16 h before collection of the supernatant.

Donor-matched iMSCs (ACS-7010; American Type Culture Collection, Manassas, VA, USA) and non-donor matched human BDMSCs (PC-500-012; American Type Culture Collection, Manassas, VA, USA) were cultured in Dulbecco’s Modified Eagle’s Medium (DMEM) [+] 4.5 g/L glucose, L-glutamine and sodium pyruvate supplemented with 10% fetal bovine serum (FBS), 1% penicillin-streptomycin and 1% non-essential amino acids in T-175 polystyrene tissue culture flasks. EV-depleted DMEM was generated via centrifugation of DMEM with supplements at 100,000 x g for 16 h before collecting the supernatant. iMSCs were passaged at ∼70% confluency for maintenance; iMSCs arrived from the manufacturer at passage 6 and were not used for EV production for functional assays after more than 10 total passages. BDMSCs were not used for EV production past 4 total passages.

Human umbilical vein endothelial cells (HUVECs) (C-12203; Promocell, Heidelberg, Germany) were cultured in T-75 tissue culture flasks coated with 0.1% gelatin using endothelial growth medium (C-C22121; PromoCell, Heidelberg, Germany). RAW264.7 mouse macrophages (T1B71; American Type Culture Collection, Manassas, VA, USA) were cultured in DMEM [+] 4.5 g/L glucose, L-glutamine and sodium pyruvate supplemented with 1% penicillin-streptomycin and 5% FBS.

THP-1 human monocytes (TIB-202; American Type Culture Collection, Manassas, VA, USA) were cultured T-175 tissue culture flasks in RPMI-1640 media supplemented with 10% heat-inactivated FBS and 1% penicillin-streptomycin inside a humidified 5% CO_2_ 37°C incubator.

THP-1 cells were maintained at a concentration between 2×10^5^ and 1×10^6^ cells/mL by passaging by dilution without centrifugation, and cells between passage 8-12 were used for the inflammatory assay.

### 2.2 EV isolation

BDMSCs or iMSCs were seeded into T-175 tissue culture flasks at ∼800,000 cells per flask and grown in EV-depleted medium. Conditioned medium was then collected over the following 3 days before being subjected to differential centrifugation steps at 1,000 x g for 10 minutes, 2,000 x g for 20 minutes and 10,000 x g for 30 minutes. The supernatant from the final centrifugation step was then passed through a 0.2 µm filter before subjection to tangential flow filtration (TFF) using a KrosFlo KR2i TFF system (Repligen; Boston, MA, USA). Using a protocol adapted from Heinemann et al., a 100-kDa MWCO MidiKros mPES membrane (D04-E100-05-N; Repligen, Boston, MA, USA) with 6 diafiltration steps and a transmembrane pressure of 5 PSI was used to concentrate samples to ∼10-15 mL^26^. Samples were then further concentrated using a 100 kDa centrifugation spin concentrator (88524; ThermoFisher Scientific; Waltham, MA, USA). Concentrated samples were then resuspended in 1x PBS and sterile filtered using a 0.2 µm syringe filter. Samples were then stored at -20^℃^ for no more than 2 weeks before use. Similarly, iPSCs were seeded into T-75s at an 8:10 dilution in colonies after passage from 6-well plates at 70% confluency. These iPSCs were grown in large particle-depleted mTESR Plus medium before media was collected and replaced daily for a total of 4 days. The collected conditioned medium was then subjected to the same differential centrifugation and TFF protocol as described above.

### 2.3 EV characterization

EV size and number was quantified via nanoparticle tracking analysis (NTA) using a NanoSight LM10 (Malvern Panalytical Limited, Malvern, UK) with version 2.3 software. Each EV sample was monitored three times with a 30 second acquisition time. Samples were diluted to achieve 20-100 particles per frame to ensure an accurate measurement with camera levels and detection thresholds kept the same between EV samples.

Transmission electron microscopy (TEM) images were obtained by using a negative stain on EV samples. Briefly, EVs were incubated with electron microscopy-grade paraformaldehyde (157-400-100; Electron Microscopy Sciences, Hatfield, PA, USA) before floating a carbon film grid (CF-200-Cu-25; Electron Microscopy Sciences, Hatfield, PA, USA) on a droplet of the EV/PFA mixture. The grids were then washed by floating on a droplet of 1x PBS before being placed on a droplet of 1% glutaraldehyde. Next, the grid was washed using a droplet of MilliQ water before being floating on a droplet of uranyl acetate replacement stain (22405; Electron Microscopy Sciences, Hatfield, PA, USA). Grids were then allowed to dry before storage and eventual imaging using a JEM 2100 LaB6 TEM (JEOL USA Incorporated; Peabody, MA, USA). Protein concentration of EV samples was determined using a bicinchoninic acid (BCA) assay using the manufacturer’s protocol (785-571; G-Biosciences, St. Louis, MO, USA). Equal amounts of protein of EV or lysate samples were then subjected to western blot analysis for ALIX (ab186429; Abcam, Cambridge, UK) at 1:1000, CD63 (25682-1-AP; ThermoFisher Scientific; Waltham, MA, USA) at 1:1000, and Calnexin (2679S, C5C9; Cell Signaling Technology Incorporated, Danvers, MA, USA) at 1:1000 overnight at 4^℃^. The following day, goat anti-rabbit IRDye 800CW (925-32210; LI-COR Incorporated, Lincoln, NE, USA) was incubated with the membrane at a 1:10,000 dilution before imaging on an Odyssey CLx imager (LI-COR Incorporated, Lincoln, NE, USA).

### 2.4 iPSC characterization

Pluripotency of iPSCs was confirmed over multiple passages and during EV production via immunofluorescence imaging. Cells were fixed using a 4% paraformaldehyde and 1% sucrose solution for 15 minutes before washing three times with 1x PBS. The cells were then permeabilized using a 6 µM magnesium chloride, 20 µM HEPES, 100 µM sodium chloride, 300 µM sucrose and 0.5% Triton-X-100 solution for 5 minutes. After additional washing with 1x PBS, cells were stained with either Oct-4 (2890S, C52G3; Cell Signaling Technology Incorporated, Danvers, MA, USA) at a 1:500, or SSEA-4 (4755S, MC813; Cell Signaling Technology Incorporated, Danvers, MA, USA) at a 1:200 dilution and incubation overnight at 4^℃^. The following day, either a goat anti-rabbit (A32731; Thermo Scientific, Waltham, MA, USA) or goat anti-mouse (A32728; ThermoFisher Scientific, Waltham, MA, USA) secondary antibody at a concentration of 10 µg/mL was incubated on the cells for 1 hour in the dark. The cells were then stained with Hoechst 33342 (62248; ThermoFisher Scientific, Waltham, MA, USA) before imaging with a Nikon Ti2 microscope (Nikon; Minato City, Tokyo, Japan).

### 2.5 Angiogenic in vitro assays

To determine endothelial gap closure, passage 5 HUVECs were seeded onto 96-well plates coated with 0.1% gelatin at a seeding density of 15,000 cells/well in endothelial growth media. After ∼24 hours, HUVECs had formed a confluent monolayer. The monolayer was then disrupted using a p200 pipette tip before washing with 1x PBS and serum-starving for 2 hours with endothelial basal media supplemented with 0.1% FBS. Following serum-starving, medium was replaced with EV treatments at a concentration of 5E9 particles/mL suspended in endothelial basal media and imaged. 16 hours later, the cells were imaged again, and the denuded area was quantified using ImageJ to determine gap closure percentage. Here, endothelial growth and basal media were used as positive and negative controls, respectively.

Tube formation assays were performed using passage 5 HUVECs. HUVECs were trypsinized and suspended in endothelial basal media supplemented with 0.1% FBS. Cells were then counted, and 75,000 cells per group were aliquoted before pelleting at 300 x g. The pelleted cells were then resuspended in their respective treatments of EVs (5E9 particles/mL) in endothelial basal medium. The resuspended HUVECs were seeded onto 24-well plates coated with growth factor-reduced Matrigel (356252; Corning, Corning, NY, USA). After 3-8 hours, phase-contrast images of tube formation were taken, and the number of branch points was determined using ImageJ.

To observe endothelial proliferation, passage 5 HUVECs were seeded onto 0.1% gelatin-coated 96-well plates at a density of 3,000 cells/well in endothelial growth media. The following day, cells were serum-starved with endothelial basal media supplemented with 0.1% FBS before replacing media with EV treatments (5E9 particles/mL) in basal media. 24 hours later, media was replaced with endothelial basal media supplemented with 0.1% FBS and CCK-8 reagent. 2-4 hours later, absorbance levels were quantified via plate reader.

### 2.6 Anti-inflammatory in vitro assays

RAW264.7 mouse macrophages were seeded into 48-well plates in DMEM supplemented with 5% FBS and 1% penicillin-streptomycin at a seeding density of 75,000 cells per well. 24 hours post-seeding, cells were pre-treated with either no treatment, 1 µg/mL dexamethasone (D4902-25MG; Sigma-Aldrich, St. Louis, MO, USA), or EV treatments (5E9 particles/mL). The following day, media was replaced with 10 ng/mL lipopolysaccharide (LPS) (L4391-1MG; Sigma-Aldrich, St. Louis, MO, USA) diluted in DMEM supplemented with 5% FBS and 1% penicillin-streptomycin for 4 hours. The conditioned media from treated RAW264.7s was then collected and stored at -80^℃^ for assessment via enzyme-linked immunosorbent assay (ELISA). After collecting the conditioned media, cells were also washed with 1x PBS and lysed in QIAzol lysis reagent (79306; QIAGEN, Hilden, Germany) for future RT-qPCR analysis.

The conditioned media from treated RAW264.7s was used to quantify levels of multiple secreted cytokines/chemokines using their respective ELISA kits including IL-6 (DY406; R&D Systems Incorporated, Minneapolis, MN, USA), TNF-⍰ (DY410; R&D Systems Incorporated, Minneapolis, MN, USA), CCL5 (DY478; R&D Systems Incorporated, Minneapolis, MN, USA), and IFN-β (DY8234; R&D Systems Incorporated, Minneapolis, MN, USA). Using the collected RAW264.7 lysate, total RNA was isolated using a RNeasy mini kit (74104; QIAGEN, Hilden, Germany) following the manufacturer’s protocol. Complementary DNA (cDNA) was then generated from total RNA samples using M-MuLV Reverse Transcriptase (M0253L; New England Biosciences, Ipswich, MA, USA) according to the manufacturer’s instructions. Following cDNA synthesis, quantitative polymerase chain reaction (qPCR) was performed using a QuantStudio 7 Flex qPCR system (4485701; ThermoFisher Scientific, Waltham, MA, USA) and PowerTrack SYBR Green Master Mix (A46109; Thermo Scientific, Waltham, MA, USA). Primer sequences used for qPCR are listed in Supplemental Table 1. The expression of mRNA transcripts was determined using a comparative Ct method normalized to either GAPDH expression and expressed as fold change of mRNA.

In “post treat” experiments looking at anti-inflammatory markers, RAW264.7 mouse macrophages were again seeded into 48-well plates in DMEM supplemented with 5% FBS and 1% penicillin-streptomycin at a seeding density of 75,000 cells per well. The following day, cells were treated with 10 ng/mL LPS for 12 hours before media was replaced with DMEM containing EV treatments (5E9 particles/mL) for 24 hours. Cells were then washed with 1x PBS and lysed using Qiazol and RNA isolation/cDNA synthesis was performed as described above for future qPCR analysis.

An NF-KB RAW264.7 alkaline phosphatase-based reporter cell line, RAWblue (raw-sp; InvivoGen, San Diego, CA, USA) was utilized to observe the relative decrease in inflammatory signaling at the transcriptional activator level. RAWblue reporter cells were plated into a 48-well plate at a seeding density of 75,000 cells per well. The following day, cells were treated with EVs (5E9 particles/mL) or their respective controls and allowed to incubate for 24 hours before stimulation with LPS (10 ng/mL) for 4 hours. After LPS stimulation, per the manufacturer’s protocol, 20 µL of conditioned media was aliquoted and mixed with Quantiblue solution (rep-qbs2; Invivogen, San Diego, CA, USA) and incubated in a 96-well plate for an additional 4 hours before quantification via plate reader.

To determine relative reactive oxygen species (ROS) concentration, a ROS assay was performed. Here, RAW264.7s were seeded into a 96-well black wall plate at a density of 12,000 cells/well. Again, cells were pre-treated for 24 hours with either EV (5E9 particles/mL) or control treatments before stimulation with LPS (100 ng/mL) for 4 hours. Post LPS stimulation, cells were washed with 1x PBS and incubated with a H2DCF2A probe (D399; ThermoFisher Scientific, Waltham, MA, USA) diluted in PBS at a concentration of 10 µM for 30 minutes. After 30 minutes, the relative fluorescence intensity was determined via plate reader.

For the THP-1 inflammatory assay, THP-1 cells were plated in 48 well plates at 150,000 cells per well with 20 nM phorbol 12-myristate 13-acetate (PMA) (P8139-1MG ; Sigma-Aldrich, St. Louis, MO, USA) supplemented in RPMI-1640 media +10% FBS and +1% penicillin-streptomycin to induce differentiation to monocyte-derived macrophages (dTHP-1), as previously described^27^. After 24 hours incubation with PMA, media was changed to fresh media and dTHP-1 cells were incubated for an additional 48 hours to allow complete differentiation. Differentiation was verified by morphological changes and adherence to tissue culture plastic. Next dTHP-1 cells were pre-treated with 2.5 μM dexamethasone as a positive control and EVs derived from iPSCs, iMSCs, and MSCs (5E9 particles/mL), and incubated for 24 hours. Then, inflammation was stimulated using 250 ng/mL LPS and 20 ng/mL IFN-γ (300-02; PeproTech, Rocky Hill, NJ, USA). Conditioned media was collected 24 hours later and stored at -80 °C until analysis of TNF-⍰ levels via ELISA (DY210; R&D Systems, Minneapolis, MN, USA).

### 2.7 EV staining and uptake

Either iPSC or iMSC-derived EVs were labeled with PKH67 (PKH67GL; Sigma-Aldrich, St. Louis, MO, USA). EVs were buffer exchanged with Diluent C using a 300 kDa MWCO Nanosep (OD300C35; Pall Corporation, New York, NY, USA) before resuspension of 200 µg of EVs in 250 µL of Diluent C. The resultant EV sample was then mixed at a 1:1 ratio with 4 µM PKH67 dye in diluent C and allowed to incubate for 5 minutes with shaking. Subsequently, 1% BSA in diluent C was added to the EV/PKH67 solution at a 1:1:1 ratio and incubated for an additional 1 minute. Dyed EV samples were then concentrated to 500 µL using a 100 kDa centrifugation concentrator. Dyed EV samples were then centrifuged at 10,000 x g for 10 minutes to remove dyed protein aggregates. To ensure removal of contaminating dye aggregates, samples were run through size exclusion columns (ICO-35; Izon, Christchurch, New Zealand) following the manufacturer’s protocol. Briefly, the first four 1 mL fractions after the void volume were collected, pooled and concentrated with a 100 kDa MWCO centrifugation concentrator before resuspension in 1x PBS and subsequent sterile filtration using 0.2 µm syringe filter. The concentration of dyed EVs was then quantified via NTA.

To assess uptake, HUVECs were seeded into endothelial growth media on 0.1% gelatin-coated 96-well black wall plates before treatment with 3E9 particles/mL in endothelial growth media 24 hours post-seeding. Cells were then washed with 1x PBS three times and either imaged using a Nikon Ti2 microscope or quantified using a plate reader. Similarly, RAW264.7s were seeded into 96-well black wall plates before treatment with 3E9 particles/mL 24 hours later. Again, cells were then washed with 1x PBS three times before either imaging or quantification via plate reader. To confirm that we were observing dyed EVs rather than uptake of dye aggregates, a mock dye solution was prepared using PBS with no EVs and subjected to the same staining and cell incubation process with both the HUVECs and RAW264.7s.

### 2.8 Animal model

24 db/db mice (40-50 g) from Jackson Laboratory (Bar Harbor, ME) were utilized for wound healing experiments. The Johns Hopkins University Animal Care and Use Committee (ACUC) approved all murine procedures, all of which followed the Johns Hopkins University ACUC Protocol (MO20M08). Briefly, mice were anesthetized with 1.5% isoflurane (Baxter Healthcare Corporation, Deerfield, IL) and the entire dorsum was shaved. An 8 mm biopsy punch (Integra, Plainsboro, NJ) was then used to wound the mice on their dorsum. On day 0, Buprenorphine Sustained-Release (1 mg/mL formulation) was locally administered subcutaneously at a dose of 0.5 mg/kg. Mice were divided into three groups, with eight mice per group: (1) Vehicle control (PBS), (2) iPSC EVs, and (3) iMSC EVs. Group matching was accomplished based on the initial wound size and animal weights on day 0. Researchers were blinded during wounding and group matching, as well as throughout the entirety of the animal experimental process. 3 days post-wounding, a total of 7.2×10^9^ EVs (determined by NTA) were injected at four quadrants intradermally into mice in the treatment groups. In each injection, there were 1.8×10^9^ EVs in a total of 50 μL of PBS. Mouse wound eschar was debrided with forceps on days 3, 6, 9, 15, and 18 to allow for clear visualization of the wound; at those timepoints, wounds were photographed and traced with clear acetate paper. Tracings were then digitized, and the wound area was quantified using ImageJ. Wound closure rates were assessed over 18 days via planimetry as the percentage of the area of the wound versus the wound size on day 3 (injection of EVs). 6 days post-wounding four mice in each group were euthanized and wounds were biopsied using a 12 mm biopsy punch. The remaining four mice were monitored, with wounds traced until day 18 where they were also euthanized, and wounds were again biopsied.

Upon wound biopsy, the tissue was cut down the center and one half was placed in RNAlater (AM7020; ThermoFisher Scientific, Waltham, MA, USA) for future RNA isolation. The other tissue half was fixed in 10% Formalin and stored overnight at 4^℃^ before briefly washing with 70% ethanol and placing in PBS before paraffin embedding and sectioning. A Leica RM2255 Motorized Rotary Microtome (Leica Biosystems; Wetzlar, Germany) was used to slice 5 µm tissue sections before mounting. H&E staining of tissue sections was then performed after deparaffinization and rehydration. Briefly, slides were incubated with hematoxylin (75810-352; VWR, Radnor, PA, USA) for 10 minutes, rinsed with running DI water, followed by a 1-minute incubation with differentiator solution (4% concentrated hydrochloric acid in 95% ethanol). Slides were then rinsed in DI water for ∼1 minute before bluing in a 1% sodium bicarbonate solution for 1 minute, washed for another ∼1 minute in DI water, placed in 95% ethanol for 1 minute, and incubated with eosin (75810-354; VWR, Radnor, PA, USA) for 1 minute and subsequently dehydrated again. Permount (SP15-100; Fisher Scientific, Hampton, NH, USA) was then added before placing cover slips on slides at a 45º angle.

For histological analysis, H&E-stained slides were scanned and digitized. To quantify wound area, which includes granulation tissue in both the dermis and new epidermis, wounds were traced and the area was measured using ImageJ using a procedure adapted from *Rhea* et al ^28^. The scar area was quantified in a similar fashion by tracing the granulation tissue within the dermis and without the inclusion of the new epidermis ^29^. For the quantification of migrating epithelial tongues, the length of new epithelium which does not yet contain dermal papillae was measured from mature epithelium (containing dermal papillae) along the wound edge to the end of the new epithelium. Again, histological analyses were performed by a blinded pathologist.

For IHC, mounted tissue sections were re-hydrated and antigen retrieval was performed by heating slides in a 10 mM Sodium Citrate buffer at 95^℃^ for 10-15 minutes. Slides were then cooled in a DI water bath, and tissue sections were circled with a liquid blocking pen. Slides were then washed with 1x TBS before blocking in a 1% bovine serum albumin (5000206; Bio-Rad, Hercules, CA, USA), and either 5% donkey (D9663-10ML; Sigma-Aldrich, St Louis, MO, USA) or goat serum (ab7481; Abcam, Cambridge, UK) solution. Slides were then incubated overnight at 4^℃^ with a 1:50 primary antibody solution of either CD206 (PA5-101657; Thermo Fisher Scientific, Waltham, MA, USA), F4/80 (MA5-16363; Thermo Fisher Scientific, Waltham MA, USA), Ly6g (14-5931-85; Thermo Fisher Scientific, Waltham, MA, USA), or CD31 (ab28364; Abcam, Cambridge UK) in a humidified chamber. Slides were then washed with 1x TBS twice for 5 minutes each and incubated with either Alexa Fluor 647 donkey anti-rabbit secondary antibody (A31573, Thermo Fisher Scientific, Waltham, MA, USA) or Alexa Fluor 647 anti-rat secondary antibody (A-21247; Thermo Fisher Scientific, Waltham, MA, USA) at a 10 µg/mL concentration for 1 hour in a dark, humidified chamber. Slides were washed with 1X TBS twice again for 5 minutes each and Vectashield Mounting Media (H-1200; Vector Laboratories, Newark, CA, USA) was added before coverslipping. Cover slips were sealed with clear fingernail polish and fluorescence images were taken on a FV3000 Laser Scanning Confocal Microscope (Olympus, Tokyo, Japan) with the same laser settings between samples at either 10x or 20x magnification over multiple fields of view per tissue section. Using ImageJ, the number of cells was determined via DAPI staining, and fluorescence intensity for the wavelength corresponding with Alexa Fluor 647 was also determined. The fluorescence intensity/number of cells was then recorded and represented as fold change over the vehicle control group.

The other half of tissue samples that were later used for RT-qPCR were incubated in RNAlater overnight at 4^℃^ before placement in a -80^℃^ freezer before RNA isolation, which occurred within ∼5 days after tissue harvesting. Using a RNeasy kit from Qiagen (74104; Qiagen, Hilden, Germany), tissue was then resuspended in Buffer RLT supplemented with β- mercaptoethanol (10 µL βME/1 mL RLT) at a ratio of 100 mg tissue to 1 mL RLT. Tissues were then homogenized with a Scilogex D160 Homogenizer (Scilogex, Rocky Hill, CT, USA) before RNA isolation using the Qiagen RNeasy kit per the manufacturer’s instructions. Reverse transcription was performed to generate cDNA in the same fashion as written above. Again, qPCR was performed in the same manner and primer sequences used for qPCR here are listed in Supplemental Table 1. The expression of mRNA transcripts was determined using a comparative Ct method normalized to β-Actin expression and expressed as fold change of mRNA.

### 2.9 Statistical Analysis

Data is presented as mean ± standard deviation. Either an ordinary one-way ANOVA was performed with Dunnett’s multiple comparisons test or a 2-sample t-tests were used to determine statistical significance. Statistical analyses were performed with Prism 9 (Graphpad Software). Statistical significance is shown as ns (p > 0.05), *p < 0.05, **p < 0.01, ***p < 0.001, or ****p < 0.0001 in figure captions.

## 3. Results

### 3.1 EV characterization and iPSC pluripotency confirmation

EVs were isolated via differential centrifugation coupled with tangential flow filtration (TFF) from the conditioned media of donor-matched iPSCs and iMSCs. Non-donor matched BDMSC EVs were also isolated in the same fashion and utilized as a benchmark/additional control in further experiments. The size distribution and concentration of each EV group was assessed via nanoparticle tracking analysis (NTA). The size distributions for each EV isolate are within the expected size ranges of EV isolates (Figure 1A). Western blots were performed on EV and lysate samples from either iMSCs or iPSCs. In these blots, EV-associated surface markers ALIX and CD63 are present in both iPSC and iMSC-derived EVs, while the cellular protein marker Calnexin is absent from EV preparations (Figure 1B). TEM images indicate that both iPSC and iMSC EVs possess the correct spherical morphology consistent with EVs (Figure 1C). To confirm pluripotency, EV producing iPSCs were stained via immunocytochemistry (ICC) for SSEA4 and OCT4 and imaged using a Nikon fluorescence microscope (Figure 1D).

**Figure 1.**
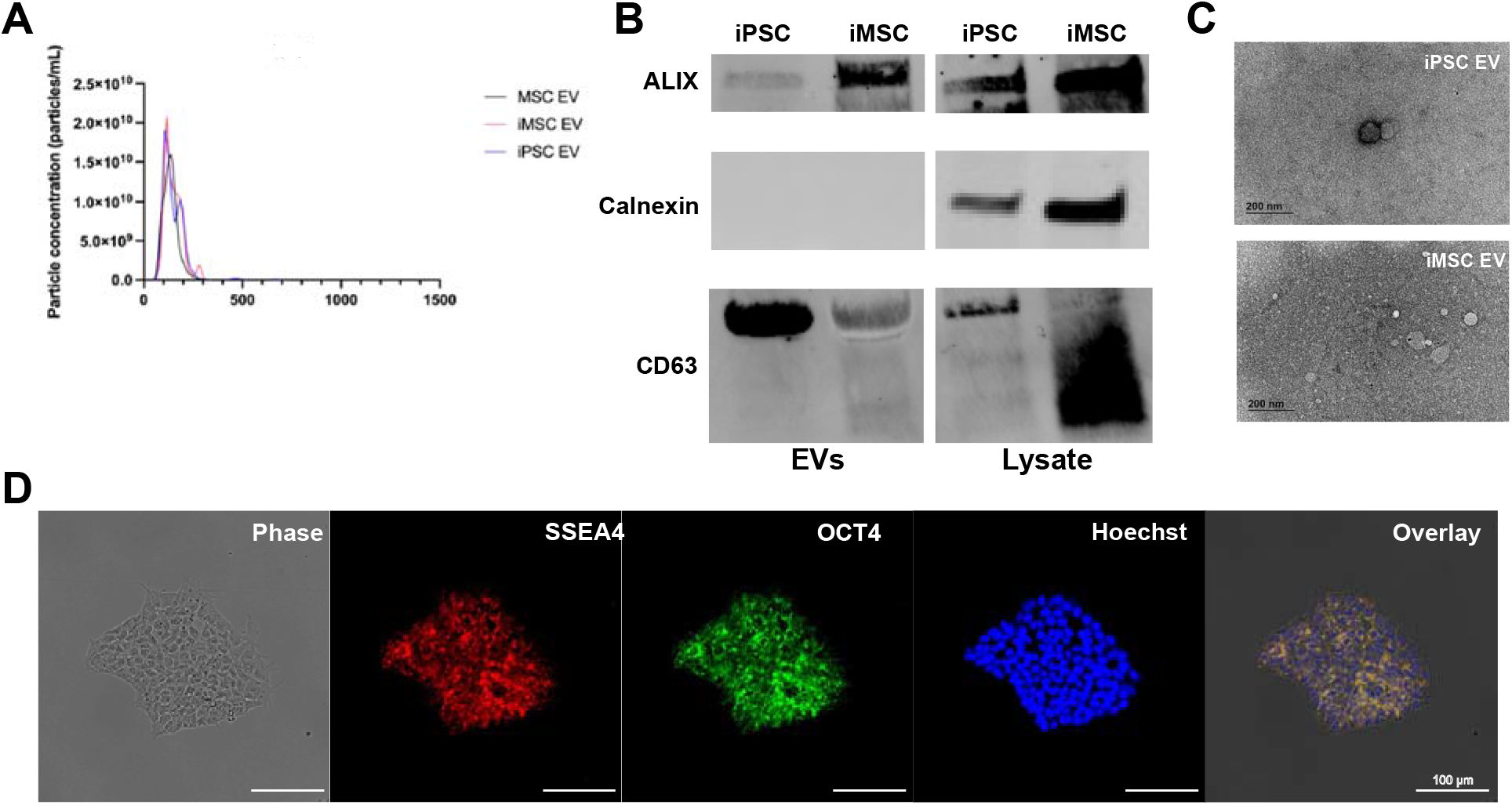
Characterization of EV size, morphology, and protein markers of EVs and parental cells. (A) NTA concentration and size distribution profiles of donor matched iPSC, iMSC and non-donor matched BDMSC EVs (B) Western blot analysis of donor-matched iPSC and iMSC EV markers ALIX and CD63 as well as Calnexin, a negative marker (C) TEM images of iPSC and iMSC EVs post-isolation (D) Representative ICC images of SSEA4 and OCT4 expression to confirm pluripotency of EV-producing iPSCs.

Confirmation of iPSC pluripotency was then performed every ∼10 passages. Meanwhile, both SSEA4 and OCT4 expression is absent from BDMSCs (acting as a control) (Supplemental Figure 1A).

Additionally, as we had observed possible particle contaminants from mTESR Plus complete media in EV isolation preparations despite being serum-free, the media was ultracentrifuged using the same protocol as employed for EV depletion to reduce possible large particle contaminants before culturing with iPSCs. The pluripotency of iPSCs cultured in this “depleted” mTESR Plus was confirmed via ICC staining for SSEA4 and OCT4 (Supplemental Figure 1B), and the depletion protocol largely removed large particle contaminants to near the lower limit of detection (Supplemental Figure 1C).

### 3.2 iPSC EVs possess similar pro-vascularization potential to donor-matched iMSC EVs in vitro

One goal of many MSC EV therapeutic approaches is to stimulate vascularization. To compare the pro-vascularization bioactivity of iPSC EVs against donor-matched iMSC EVs, a tube formation assay was performed with HUVECs grown on growth-factor reduced Matrigel. At a dose of 5×10^9^ particles/mL as assessed by NTA, donor-matched iMSC and iPSC EVs produced endothelial tube-like structures with similar amounts of branch points per field of view, which is significantly more compared to untreated HUVECs in endothelial basal media (Figure 2A). Additionally, a gap closure assay was performed on a confluent monolayer of HUVECs and again treatment with either iMSC or iPSC EVs yielded similar pro-vascularization potential over basal media control (Figure 2B).

**Figure 2.**
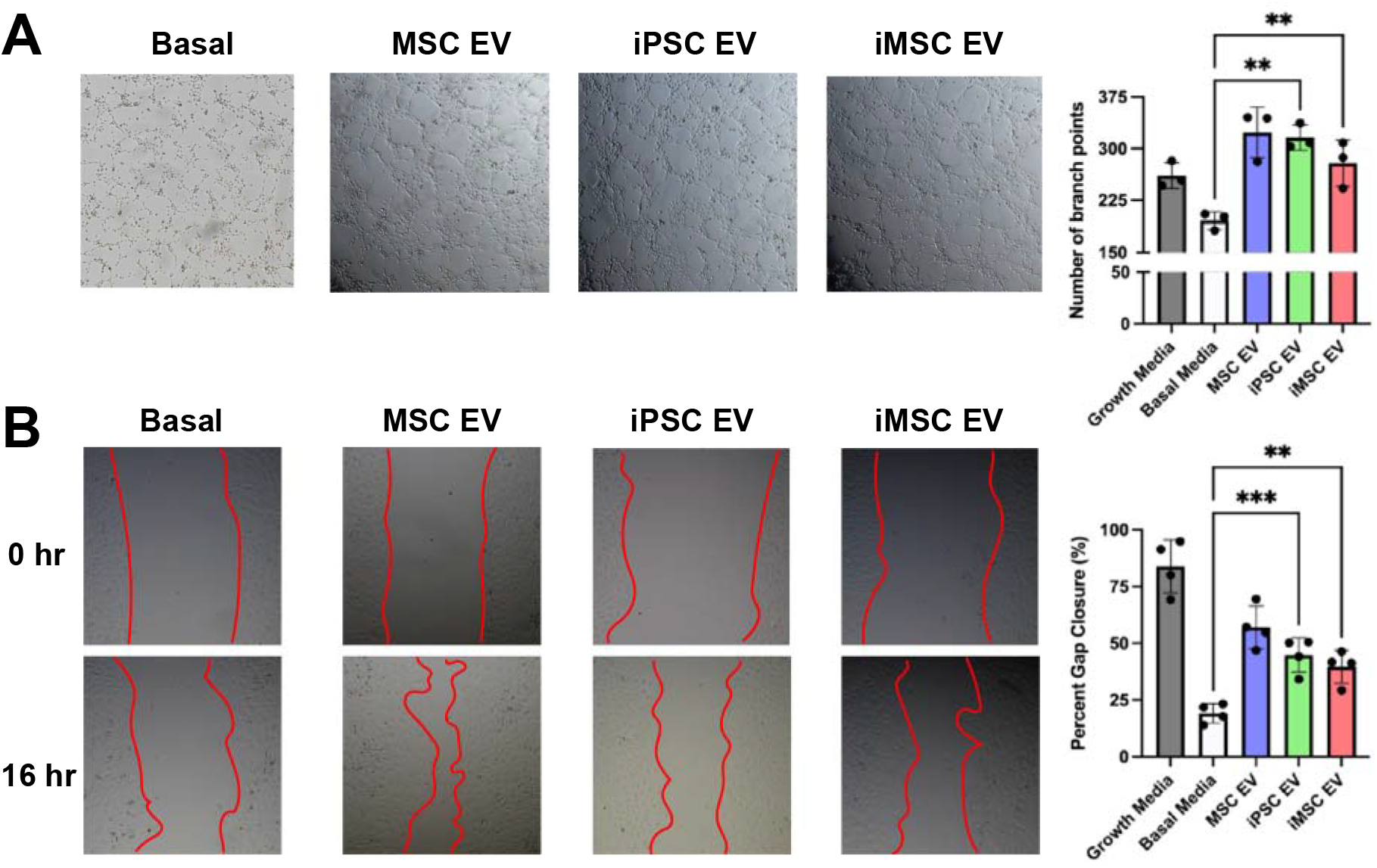
iPSC EVs have similar pro-angiogenic potential to donor-matched iMSC EVs. (A) After resuspension in EV treatments, HUVEC tube formation was quantified by the number of branch points per bright field image (n=3). (B) After inducing a scratch, HUVECs were treated with EVs in basal media and the percentage of gap closure was assessed using bright field images (n=4). All values were expressed as mean ± standard deviation (**p < 0.01, ***p < 0.001)

To assess the ability of HUVECs to take up iPSC and iMSC EVs *in vitro*, EVs as well as a PBS mock control were exposed to fluorescent PKH67 dye and subjected to SEC to remove unbound dye before culturing with HUVECs for 24 hours. HUVECs were then washed and imaged on a Nikon fluorescence microscope or fluorescence intensity was quantified via plate reader, with the data indicating similar uptake levels in HUVECs for both iPSC and iMSC EVs (Supplemental Figure 2A). Additionally, EV-mediated HUVEC proliferation was assessed using a CCK8 assay. At a dose of 5×10^9^ particles/mL, iPSC EVs induced proliferative bioactivity in HUVECs *in vitro*, whereas donor-matched iMSC EVs at the same dose did not (Supplemental Figure 2B).

### 3.3 iPSC EVs exhibit similar to superior anti-inflammatory bioactivity when compared to donor-matched iMSC EVs

As MSC EVs have been extensively reported to possess anti-inflammatory properties, an *in vitro* LPS-stimulated mouse macrophage model was used to benchmark the anti-inflammatory properties of IPSC EVs against donor-matched iMSC EVs ^30^. iPSC EV treatment significantly reduced the secretion of the pro-inflammatory cytokines/chemokines IL-6, TNF-⍰, CCL5, and IFN-β compared to controls (Figure 3A). iMSC EVs reduced IL-6, CCL5, and IFN-β levels compared to controls, but not TNF-⍰, and in each case the reduction was less than what was achieved by donor-matched iPSC EVs, with the disparities for TNF-⍰ and CCL5 being statistically significant (Figure 3A). EV uptake by RAW264.7 cells was confirmed (Supplemental Figure 3), and validation of the dose-dependent nature of the anti-inflammatory effect of iPSC EVs was carried out for IL-6 (Figure 3B). Additionally, the ability of iPSC EVs to reduce inflammatory IL-6 levels did not change with increased passage (Figure 3C), supporting the concept that iPSCs can serve as sources for reproducible and scalable biomanufacturing of therapeutic EVs, in contrast to donor-sourced primary MSCs ^13^. RT-qPCR analysis on the cell lysate revealed that mRNA expression of IL-6, as well as TNF-⍰ and iNOS, were also significantly reduced by either iPSC or iMSC EV treatment (Supplemental Figure 4A). Finally, a RAW264.7 NF-KB reporter cell line was used to determine the ability of EVs to modulate inflammatory activation at the transcriptional activator level; both iPSC and iMSC EVs reduced NF-KB activity compared to control as measured by alkaline phosphatase secretion (Supplemental Figure 4B).

**Figure 3.**
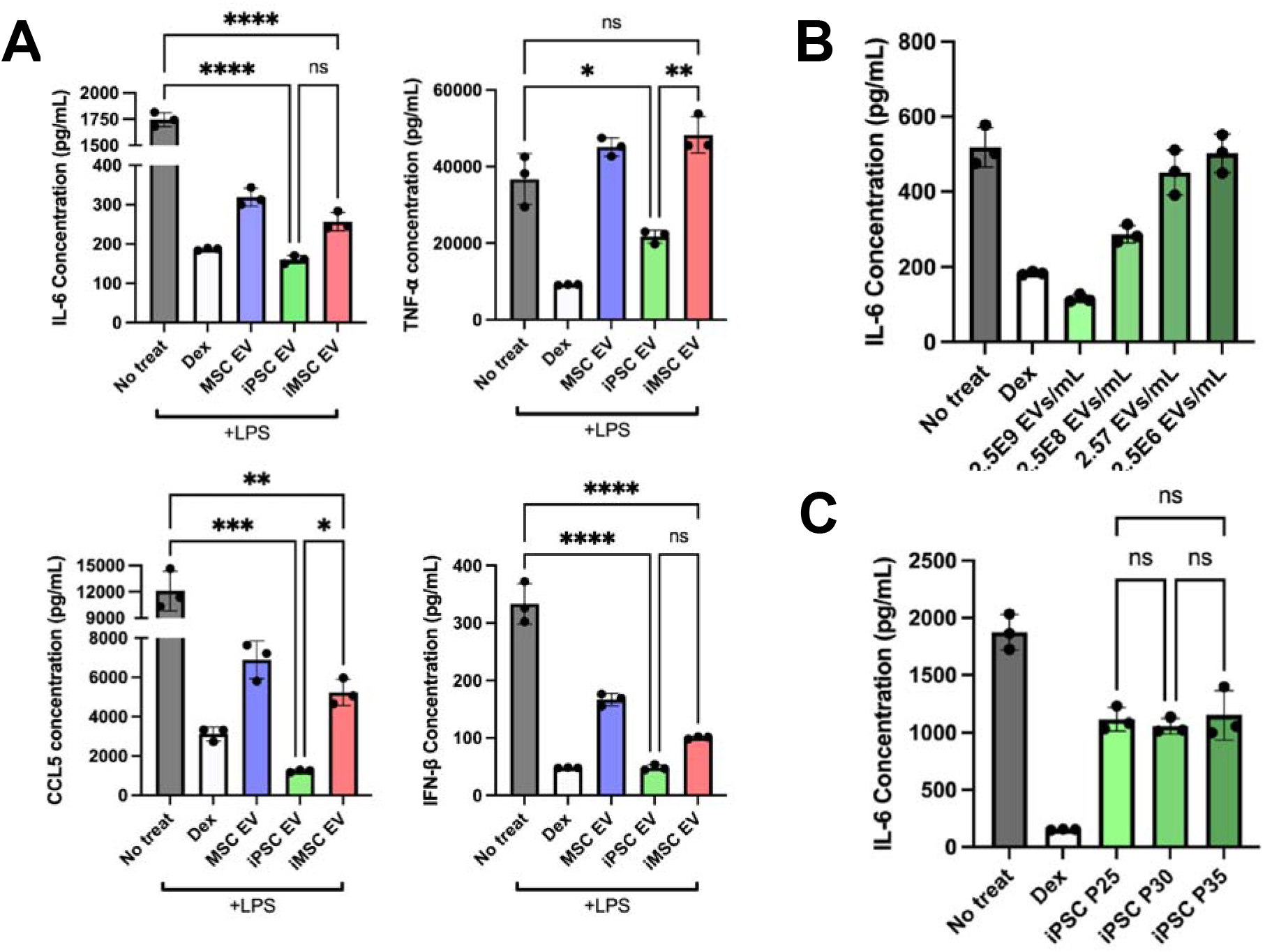
iPSC EVs possess superior anti-inflammatory properties compared to donor-matched iMSC EVs. (A) RAW264.7 mouse macrophages were pre-treated with the indicated EV treatments before LPS stimulation. The cell supernatant was then collected and IL-6, TNF-⍰, CCL5, and IFN-β protein levels were quantified using ELISA (n=3). (B) RAW264.7 mouse macrophages were pre-treated with iPSC EVs at the indicated doses before LPS stimulation (10 ng/mL). Cell supernatants were collected and IL-6 levels were quantified using ELISA (n=3). (C) EVs isolated from iPSCs over multiple passages were used in the same LPS-stimulated RAW264.7 macrophage assay and IL-6 levels in the cell culture media was quantified via ELISA (n=3). All values were expressed as mean ± standard deviation (*p < 0.05, **p < 0.01, ***p < 0.001, ****p <0.0001).

Next, the potential of iPSC EVs to induce cellular changes related to inflammation resolution and repair was examined. As resolution typically occurs after an initial acute inflammation response, a “post-treat” cellular model was employed, where RAW264.7 macrophages were stimulated with 10 ng/mL LPS for 12 hours before treatment with EVs at a dose of 5×10^9^ particles/mL for 24 hours ^31^. Via RT-qPCR analysis, expression of the anti-inflammatory cytokine IL-10 and the “M2” macrophage marker CD206 were both increased by treatment with iPSC EVs, while iMSC EVs had smaller effects (Figure 4A). Another key mechanism in inflammation resolution is the ability to dampen the release of reactive oxygen species (ROS), which have been established to be a partial driver of inflammatory responses in injury ^32^. Thus, EV-pre-treated RAW264.7s were stimulated with LPS (100 ng/mL) and H2DCFDA fluorescent probe was subsequently added to quantify relative ROS levels. Treatment with either iPSC EVs or IMSC EVs reduced fluorescent signals compared to LPS-stimulated RAW264.7s that received no pre-treatment (Figure 4B).

**Figure 4.**
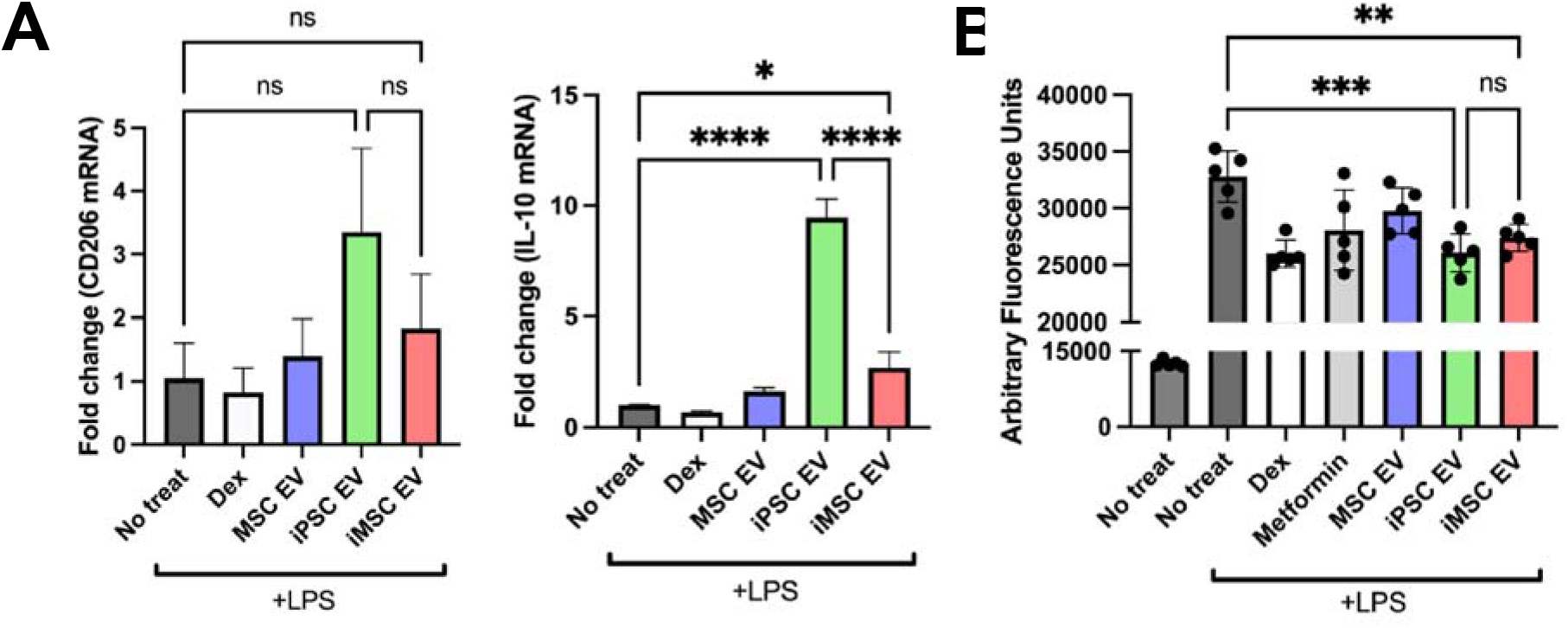
iPSCs EVs resolve inflammation by transitioning macrophages to an “M2” phenotype and reduce ROS levels. (A) In a “post-treat” regime, where RAW264.7s were stimulated with LPS, treated with EVs before lysis, anti-inflammatory macrophage markers/cytokine mRNA expression levels were quantified via RT-qPCR (n=3). (B) RAW264.7 mouse macrophages were pre-treated with EVs before LPS stimulation (100 ng/mL) and subsequent ROS quantification using a H2DCFDA fluorescent dye along with fluorescence quantification via plate reader (n=6). All values were expressed as mean ± standard deviation (*p < 0.05, **p < 0.01, ***p < 0.001), ****p < 0.0001).

To confirm that iPSC EV preparations were effective in reducing inflammatory phenotypes in human cells in addition to mouse macrophages, a TNF-⍰ stimulated HUVEC assay was used to assess expression of adhesion molecules utilized by leukocytes for extravasation into local sites of inflammation. iPSC EV treatment led to marginally decreased expression of VCAM-1 in this model (Figure 5A). Additionally, utilizing a stimulated human THP-1 assay, both iPSC EVs and iMSC EVs induced a robust decrease in TNF-⍰ secretion as assessed by ELISA (Figure 5B).

**Figure 5.**
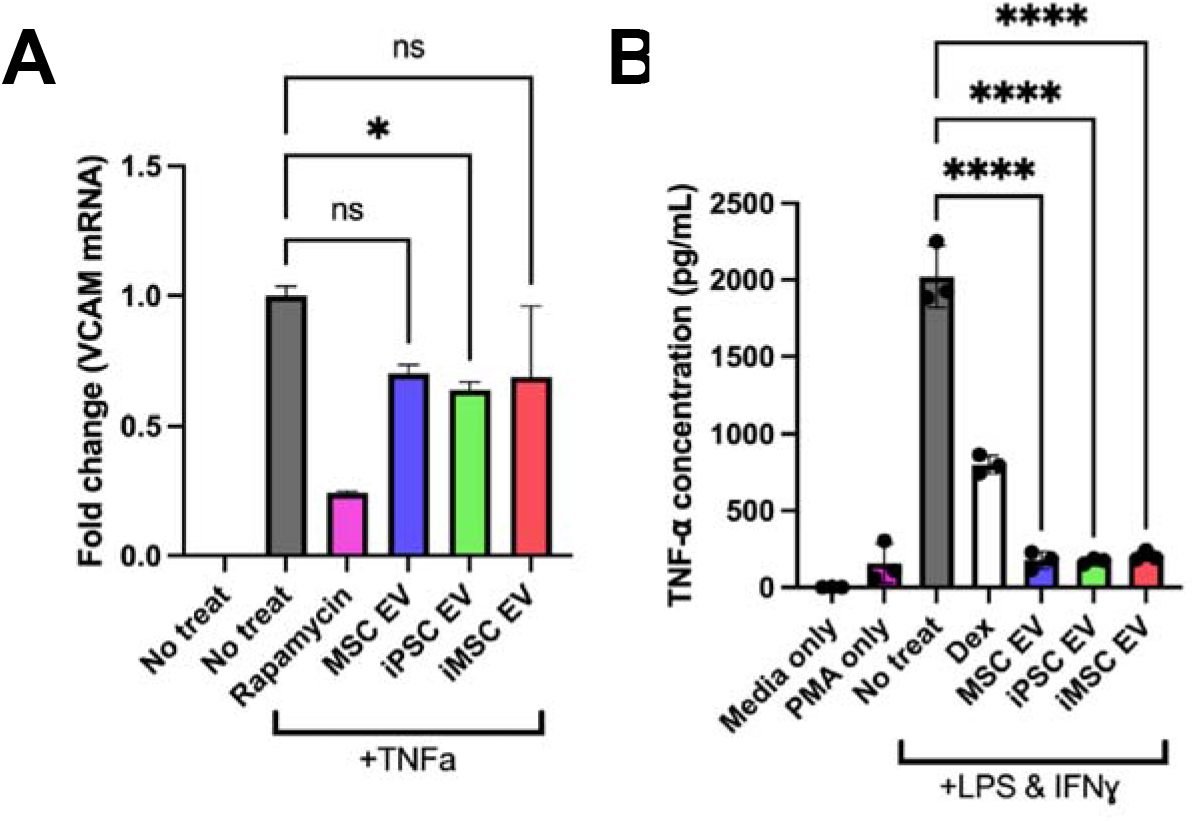
iPSC and iMSC EVs have anti-inflammatory effects in human cell models. (A) HUVECs were pre-treated with EVs for 24 hours at a dose of 5E9 particles/mL before stimulation with 10 ng/mL TNF⍰ for 16 hours before lysing and quantification the endothelial adhesion marker, VCAM1 via RT-qPCR. (B) We looked to confirm the anti-inflammatory effects of our EV samples in a human LPS-stimulated THP1 macrophage assay. The conditioned media of stimulated THP1s was collected and TNF-⍰ levels were quantified via ELISA. All values were expressed as mean ± standard deviation (*p < 0.05, ****p < 0.0001).

Our group previously reported on media contaminants affecting the outcomes of anti-inflammatory assays involving EVs ^33^. To verify that the reductions in pro-inflammatory cytokine secretion in this model were due to EVs and not media contaminants, the RAW264.7 pre-treat assay was performed using mTESR Plus that had undergone the EV isolation process. We observed that the mTESR Plus depletion protocol was effective at removing contaminants that may skew anti-inflammatory assay results; additionally, we saw that upon culture with iPSC EVs, the anti-inflammatory effect was restored (Supplemental Figure 5A). Another concern was the possibility that iPSC EV treatment was toxic to the RAW264.7s in this assay, leading to lower cytokine levels. However, using a CCK8 assay, we observed that iPSC EV treatment actually increased cell viability and number (Supplemental Figure 5B).

### 3.4 iPSC EVs reduce gross wound size in a db/db mouse wound healing model

To compare the anti-inflammatory and pro-angiogenic properties of iPSC EVs and iMSC EVs in a more rigorous setting, a wound healing model in db/db mice was utilized (Figure 6A). Wounds were traced every three days after EV injection to monitor wound size/closure over time. However, no significant increase in wound closure rate induced by either iPSC EVs or iMSC EVs was observed (Figure 6B). This was not surprising, as the wound healing model did not employ stenting, and thus wound closure was likely driven by the contraction of the surrounding skin tissue rather than the growth of new epithelial tissue ^34^. For a more relevant assessment of healing in this model, wound area was examined histologically. Blinded analysis of H&E-stained tissue slices from skin collected 18 days after initial wounding indicated an ∼45% reduction in total wound area in iPSC EV-treated mice compared to the PBS control, while iMSC EV treatment had no effect (Figure 6C). To confirm these findings, the lengths of wounds were measured by tracing the outer wound edges. Again, iMSC EV treatment was shown to have little effect in reducing wound length, while iPSC EV treatment induced a non-statistically significant ∼25% decrease in wound length (Figure 6D). Further, a significant ∼50% reduction in scar area was associated with iPSC EV treatment, with a non-significant 10% reduction achieved via iMSC EV treatment when compared to the vehicle control (Supplemental Figure 6B).

**Figure 6.**
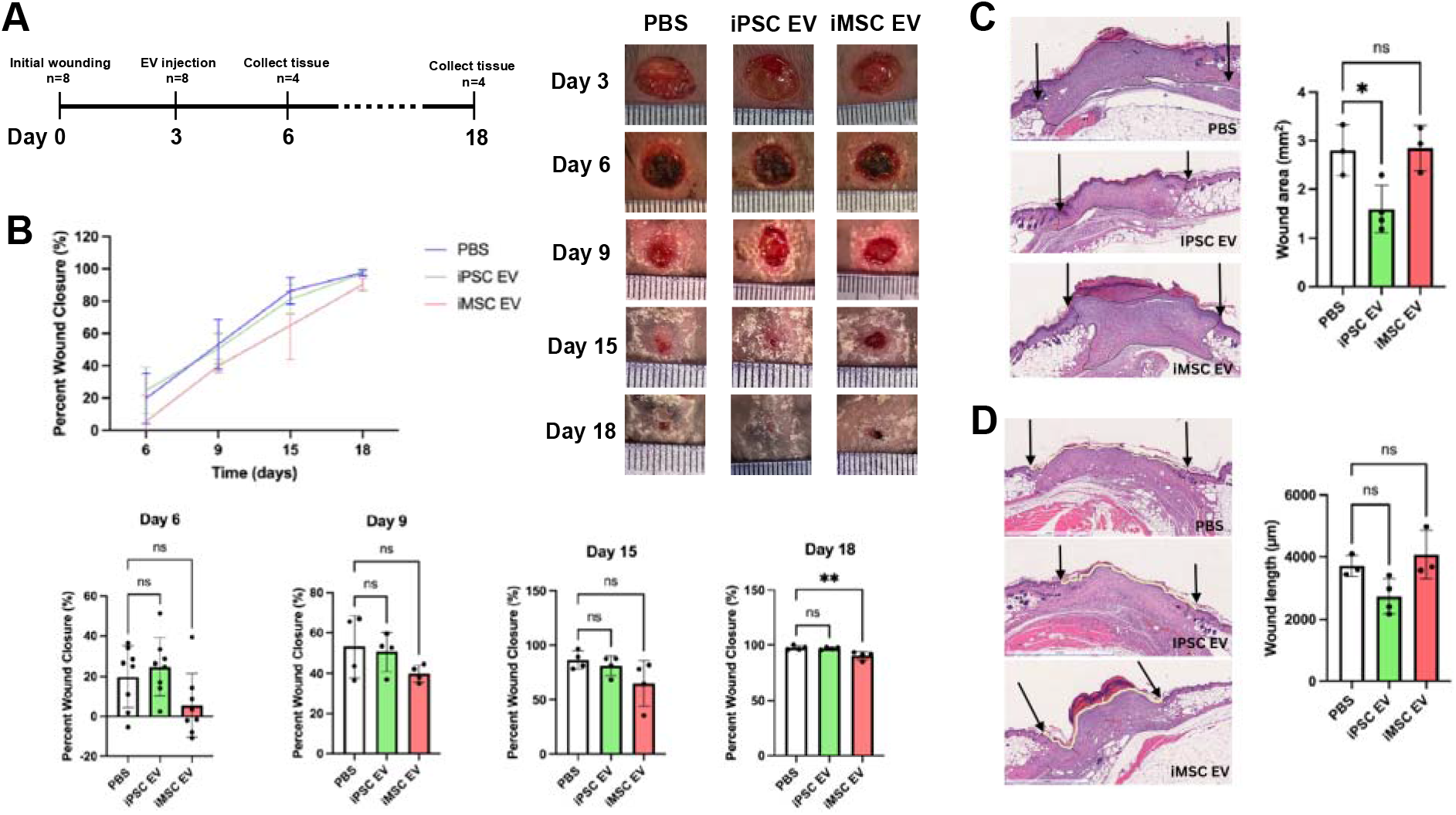
iPSC EVs improve wound tissue architecture during healing in a db/db mouse wound model. (A) Timeline of wounding, injection and tissue harvesting. (B) Wound closure rate was assessed over 15 days via planimetry from representative wound images for wounds treated with donor-matched iPSC and iMSC EVs as well as a PBS vehicle control (n=4-8). (C) Representative images of H&E-stained wound beds 18 days post-wounding. Total wound area was quantified by tracing the granulation tissue within the wound bed (n=3-4) (D) Representative images of H&E-stained wound beds 18 days after wounding. Wound length was quantified by tracing and measuring the outer wound edge (n=3-4). All values were expressed as mean ± standard deviation (*p < 0.05, **p < 0.01)

### 3.5 iPSC EVs induce anti-inflammatory macrophage phenotypes in vivo

To assess potential mechanisms of iPSC EV wound repair effects, anti-inflammatory activity was investigated following from the results of the prior *in vitro* experiments. At a timepoint reported to coincide with the inflammation resolution phase of wound healing (6 days after initial wounding) ^35^, mice were sacrificed and a punch biopsy of the wound area was taken and processed for histological and immunohistochemical analyses. Images of H&E-stained tissues revealed significant amounts of necrotic tissue near the wound surface, underlying fibrotic tissue, and leukocyte infiltrate (Figure 7A), the latter of which can be instrumental to either resolution or persistence of chronic wounds ^36^. Additionally, re-epithelization of the wound bed occurred at an enhanced rate in iPSC EV-treated mice, as evidenced by an ∼85% increase in new epithelial tongue length over the PBS control, while iMSC EV-treated wounds were not significantly different than vehicle-treated wounds (Figure 7A). Immunohistochemistry (IHC) for F4/80, a general macrophage marker ^36^, indicated an ∼30% increase in total macrophage infiltration in iPSC EV-treated wounds compared to vehicle-treated mice, with no significant increase over PBS control with iMSC EV treatment (Figure 7B).

**Figure 7.**
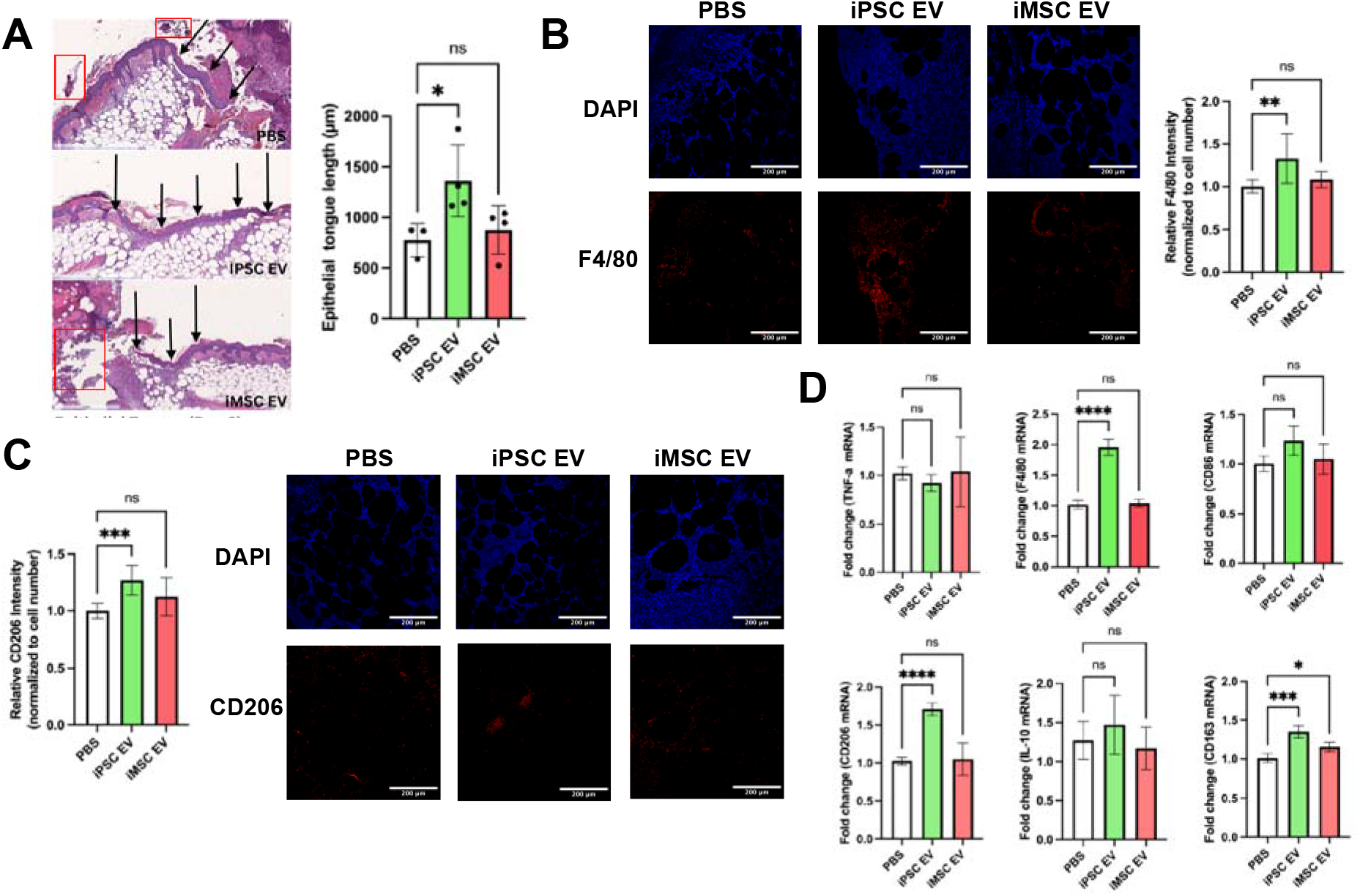
Inflammation-resolving macrophages are increased upon iPSC EV treatment. (A) Representative images of H&E-stained wound beds 6 days post-wounding. Necrotic and apoptotic tissue are highlighted with red boxes. New epithelium was measured in length from the mature epithelium along the wound edge demarcated by black arrows. (n=3-4) (B) Images of F4/80 IHC-stained tissues 6 days after wounding. Total F4/80 fluorescence intensity was quantified and normalized to cell number via DAPI over multiple fields of view. (n=4) (C) Representative images of CD206 IHC-stained tissues 6 days post-wounding. Again, CD206 fluorescence intensity was normalized to cell number for quantification. (n=4) (D) Inflammatory/macrophage cytokine and surface markers were quantified via RT-qPCR of mRNA isolated from bulk wound bed tissue 6 days post-wounding (n=4). All values were expressed as mean ± standard deviation (*p < 0.05, **p < 0.01, ***p < 0.001, ****p < 0.0001)

To determine whether these infiltrating macrophages participated in inflammation persistence or resolution, IHC assessment of CD206, a “M2” macrophage marker indicative of macrophages that actively aid the repair process ^37^, was performed. An ∼25% increase in CD206 intensity in iPSC EV treated wounds over the PBS control was detected, while iMSC EVs did not induce a significant increase (Figure 7C). While macrophages are critical to the wound healing process, they are not the sole driver of inflammation persistence/resolution, as neutrophils are another key leukocyte that drives the initial inflammatory response in wounds ^38^. Thus, IHC for Ly6G expression was performed to assess neutrophil infiltration (as well as monocytes/granulocytes) ^39^. A non-statistically significant decrease (∼30%) in intensity was observed for iPSC EV treated wounds, while a significant (∼50%) increase in Ly6G intensity was associated with iMSC EV treatment compared to PBS control (Supplemental Figure 7A), indicative of a potential disparity in the mechanisms of action of iPSC EVs and iMSC EVs. To validate IHC findings, bulk RNA isolation from the wound bed tissue was performed before RT-qPCR for pro-inflammatory markers TNF-⍰ and iNOS, activated macrophage marker CD86, and anti-inflammatory markers/cytokines CD206, IL-10, CD163, and TGF-β (Figure 7D, Supplemental Figure 7B) ^36^. No significant decreases in pro-inflammatory TNF-⍰ or iNOS were detected via this method with iPSC EV treatment compared to the PBS control; additionally, IL-6 levels were too low to quantify using RT-qPCR (data not shown). Surprisingly, iNOS expression was increased with iMSC EV treatment (Supplemental Figure 7B). However, when looking at expression of anti-inflammatory “M2” macrophage markers^36^, a robust ∼70% increase in CD206 along with an ∼35% increase in CD163 expression were observed associated with iPSC EV treated wounds (Figure 7D). Additionally, a non-significant increase in IL-10 and TGF-β expression was observed in iPSC EV-treated wounds (Figure 7D, Supplemental Figure 7B). Interestingly, the changes in anti-inflammatory markers/cytokines for iMSC EV treated wounds were all relatively marginal, indicating a muted immunomodulatory overall effect.

While it was expected that resolution of inflammation would largely be complete by 18 days post-wounding, F4/80 IHC and fluorescence imaging was performed at this time point for confirmation. As expected, macrophage infiltration was low and unchanged between the PBS control and both iPSC EV- and iMSC EV-treated wounds, indicating the inflammatory response was largely resolved by this time point (Supplemental Figure 7C). CD206 IHC was also performed at this time point, with results showing similar normalized intensity between PBS, iPSC EV-, and iMSC EV-treated wounds, with a slight ∼15% increase in CD206 intensity in the iMSC EV group (Supplemental Figure 7D). This may indicate that iMSC EV treatment of wounds induces persistence of tissue repair-associated macrophages. Alternatively, when considering the CD206 data for wounds treated with iMSC EVs from the previous timepoint (Figure 7C,D), it could be that this tissue resolving effect was simply delayed compared to the iPSC EV and PBS groups.

### 3.6 iPSC EVs marginally increase re-vascularization in a db/db mouse wound healing model

As the wound should progress towards to the proliferative phase of repair by 18 days post-wounding, where re-vascularization plays a critical role in supplying nutrients to the repaired tissue, blood vessel formation was assessed at this timepoint via CD31 IHC ^35^. Non-statistically significant ∼35% and ∼20% increases in CD31+ staining were associated with iPSC EV and iMSC EV treatments, respectively (Figure 8A). Bulk RNA isolation from the wound tissue was again performed, this time to evaluate expression of pro-angiogenic markers VEGF, FGF1, FGF2, Angiopoietin2, PDGFb, and IGF1 via RT-qPCR. Overall, no changes in VEGF, PDGFb, FGF1, FGF2, or Angiopoietin2 expression were observed with either iPSC EV or iMSC EV treatment (Figure 8B). Interestingly, there was an ∼40% increase in IGF1 expression in iPSC EV-treated wounds compared to the vehicle control (Figure 8B). Overall, this lack of evidence for substantial increases in vascularization is not entirely surprising, as we have previously demonstrated that unmodified MSC EVs had only marginal effects in increasing the number of blood vessels in the same wound healing model ^40^. However, it is important to note that iPSC EVs were similarly ineffective at substantially improving angiogenesis in this wound model when compared to donor-matched iMSC EVs.

**Figure 8.**
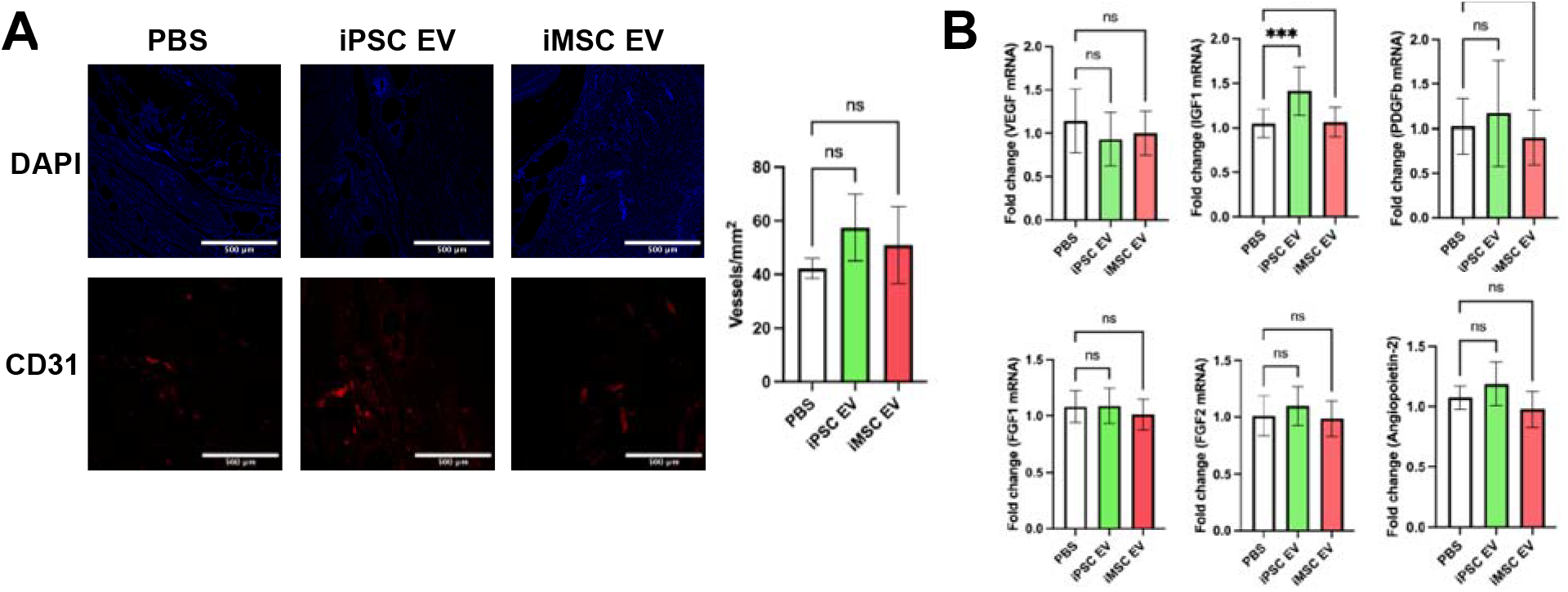
iPSC EVs marginally affect re-vascularization during the proliferative phase of wound healing. (A) Representative images of CD31 immunochemistry-stained tissue. Blood vessels were counted within in a 1mm^2^ field of view. (n=4) (B) Pro-angiogenic growth factor expression was quantified via RT-qPCR from bulk mRNA isolated from wound tissue 18 days after wounding (n=4). All values were expressed as mean ± standard deviation (***p < 0.001).

## 4. Discussion

Clinical trials and investment from industry into EV-based therapeutics continue to increase, driving the need to address the barriers that remain to their ultimate clinical translation ^41, 42^. One key challenge is the issue of scalability. In particular, MSCs, a popular cell type for therapeutic EV production, have been widely reported to possess limited expansion capabilities *ex vivo*, thus capping the number of cells and EVs that can be produced from a singular MSC line ^10^. The effects of donor variability on MSC function are well reported, and this translates to their EVs as well, further limiting reproducible production of EVs with predictable therapeutic characteristics ^16, 43^. Additionally, our group has previously demonstrated that EVs isolated from MSCs at higher passages begin to possess dampened functional bioactivity, imposing yet another limitation on the number of viable therapeutic EVs that can be obtained ^13^. Therefore, validation of scalable sources for therapeutic EV production is crucial to the continued development of this class of therapeutics.

The use of iPSCs for EV production is thus compelling due to their self-renewing capabilities ^44^. EVs from iPSCs have been investigated in several applications to date, with promising results in models of cardiac injury, liver fibrosis, and cellular aging ^23-25, 45^. The results of this study expand into the application space of wound repair, with surprising implications related to differentiation of iPSCs for EV production. Based on previous reports that iPSC EVs possess pro-angiogenic properties *in vitro*, we expected undifferentiated iPSC EVs to perform similarly to iMSC EVs in an *in vitro* angiogenic screen, which was born out in the results (Figure 2) ^24, 45^. However, due to the well-established anti-inflammatory properties of MSC EVs, we also expected iMSC EVs to outperform iPSC EVs in anti-inflammatory assays ^46^. Yet, we observed that iPSC EVs may have superior anti-inflammatory properties to EVs from donor-matched iMSCs in terms of both reducing pro-inflammatory phenotypes and inducing anti-inflammatory/inflammation resolving phenotypes (Figures 3-5). This finding is critical, as it further bolsters the rationale behind using EVs from undifferentiated iPSCs over iMSCs in addition to the production advantages inherent in avoiding additional differentiation steps. Additionally, we found that iPSC EVs retain bioactivity over many passages of the producer cells (Figure 4C), further emphasizing their enhanced utility compared to donor MSC EVs with respect to scalability ^13^.

Given that wound healing is a complex process that involves re-vascularization as well as macrophages playing an active role in both the promotion and resolution of inflammation, we hypothesized that this application may be appropriate for iPSC EVs ^47^. While there was no increase in wound closure overall, this may be significantly attributed to limitations of the chosen model – wound closure in mouse wound healing models is affected not only by re-epithelization, as is the case in human wound healing, but also the contraction of surrounding skin tissue, which is the critical driver of wound closure in mouse models (Figure 6) ^34^. Meanwhile, histological analysis demonstrated that iPSC EV treatment did increase the rate of new epithelium formation at earlier time points (Figure 7A), as well as smaller overall wound area and length at later time points (Figure 6C,D). Based on the slight increase in wound closure at the first time point associated with iPSC EV injection, the dosing scheme could be modified either by increasing the bolus dose or by employing repeated doses.

As there are distinct limitations with respect to wound closure rate in our model, we also assessed some of the cellular and molecular responses within the wound bed. We did not observe a decrease in pro-inflammatory mRNA expression levels upon EV treatment, which was surprising after observing such robust decreases in our *in vitro* model (Figure 3, Supplemental Figure 4). It is possible that by harvesting tissues 3 days post-injection (6 days post-wounding), the peak wound inflammatory phase, which typically occurs 3-4 days post-wounding ^35^, was missed. This also applies with respect to peak neutrophil infiltration. However, we did observe an increase in macrophage infiltration within wounds 3 days after iPSC EV treatment as assessed by F4/80 IHC (Figure 7B). These infiltrating activated macrophages are likely “M2” macrophages, or inflammation resolution/tissue repair macrophages, as indicated by higher expression levels of CD163 and CD206 in iPSC EV-treated groups (Figure 7C,D)^37^. Due to these findings, we also looked at whether anti-inflammatory cytokines such as IL-10 and TGFβ mRNA expression levels were increased, but saw only marginal, non-significant increases with iPSC EV treatment (Figure 7D, Supplemental Figure 4B). Interestingly, we did not observe many differences in inflammatory markers with iMSC EV treatment and actually observed increased iNOS expression and Ly6G intensity (Figure 7D, Supplemental Figure 4A), indicative of a discordance between the *in vitro* and *in vivo* results. These contradicting results may be due to the timing of *in vivo* sample acquisition as well as other factors.

As wounds begin to move into the proliferative phase of the healing process, nutrient supply to the repaired tissue is critical to inducing an environment hospitable to repair where promotion of angiogenesis is key ^48^. Thus, 15 days post-injection (18 days post-wounding), we assessed whether EV treatment improved re-vascularization of the tissue^35^. Overall, no significant changes in promotion of angiogenesis in the wounds were observed associated with either iMSC or iPSC EV injection compared to vehicle control (Figure 8). These results aren’t surprising given that we previously reported that unmodified MSC EVs had little effect in increasing blood vessel number in the same model ^40^. Further, in addition to no significant changes in blood vessel density, there was little difference in pro-angiogenic mRNA expression within the wound bed between EV treatments and the vehicle control (Figure 8B). However, it was observed that IGF1 was increased in iPSC EV treated wounds by ∼40% (Figure 8B). This is particularly interesting in a diabetic wound healing model specifically, given the role of IGF1 in insulin regulation ^49^. This result may also be supported by a recent study that profiled cargos within iPSC EVs, showing that many of these cargos are involved with modulating metabolism and aging, which IGF1 is also involved in ^50, 51^.

Given evidence that macrophage phenotypes are also related to cellular metabolism, it is possible that iPSC EVs may impart the observed “M2” transition through a similar pathway ^52-54^. However, further studies into the mechanism behind these anti-inflammatory phenomena are needed. Mechanistic studies are also needed to understand and rationally design enhanced iPSC EV-based therapeutics in the future. Lastly, development of downstream processes to sustain scalable production, such as utilization of bioreactors or reducing the cost of media formulations, would aid in the translation of iPSC EVs to the clinic ^8, 55^. Despite the bevy of possible studies that remain, the results here further support the use of iPSC EV-based therapeutics by demonstrating that iPSCs may be a superior alternative therapeutic EV source to iMSCs with respect to both therapeutic efficacy and scalability.

## Supporting Information

Supporting Information is available online.

## Acknowledgements

The authors acknowledge the University of Maryland School of Medicine’s Pathology Histology Core – Baltimore, Maryland for consultation on services provided. This work was supported by the National Institutes of Health (HL141611, NS110637, GM130923, HL141922, HL159590 to SMJ; HL007698 to EP) and the National Science Foundation (1750542 to SMJ). DL and TS were supported by A. James Clark Doctoral Fellowships from the University of Maryland. NJP was supported by a MPower Graduate Fellowship from the University of Maryland. JWH was supported by the NIH (HL141611) and the Hendrix Burn/Wound Fund of Johns Hopkins University.

