## Supplemental Material for "Induced pluripotent stem cell-derived extracellular vesicles promote wound repair in a diabetic mouse model via an anti-inflammatory immunomodulatory mechanism"

**SUPPLEMENTAL INFORMATION**

***Supplemental Table 1.*** *Primer sequences used for RT-qPCR*

| *β-actin* | FWD  REV | CAGGTCATCACTATTGGCAA  AGGTCTTTACGGATGTCAAC |
| --- | --- | --- |
| *GAPDH* | FWD  REV | GTGTTCCTACCCCCAATGTG  CATCGAAGGTGGAAGAGTGG |
| *TNF-⍺* | FWD  REV | GCATGATCCGAGATGTGGAACTGG  CGCCACGAGCAGGAATGAGAAG |
| *iNOS* | FWD  REV | ACTCAGCCAAGCCCTCACCTAC  TCCAATCTCTGCCTATCCGTCTCG |
| *IL-6* | FWD  REV | AATTTCCTCTGGTCTTCTGGAGT  GTGACTCCAGCTTATCTCTTGGT |
| *CD86* | FWD  REV | ACGGAGTCAATGAAGATTTCCT  GATTCGGCTTCTTGTGACATAC |
| *CD206* | FWD  REV | CCTATGAAAATTGGGCTTACGG  CTGACAAATCCAGTTGTTGAGG |
| *IL-10* | FWD  REV | TTCTTTCAAACAAAGGACCAGC  GCAACCCAAGTAACCCTTAAAG |
| *CD163* | FWD  REV | GGCTAGACGAAGTCATCTGCAC  CTTCGTTGGTCAGCCTCAGAGA |
| *TGF-β* | FWD  REV | TGATACGCCTGAGTGGCTGTCT  CACAAGAGCAGTGAGCGCTGAA |
| *IL-4* | FWD  REV | ATCATCGGCATTTTGAACGAGGTC  ACCTTGGAAGCCCTACAGACGA |
| *IL-13* | FWD  REV | AACGGCAGCATGGTATGGAGTG  TGGGTCCTGTAGATGGCATTGC |
| *VEGF* | FWD  REV | CTGCTGTAACGATGAAGCCCTG  GCTGTAGGAAGCTCATCTCTCC |
| *IGF-1* | FWD  REV | GTGGATGCTCTTCAGTTCGTGTG  TCCAGTCTCCTCAGATCACAGC |
| *PDGFb* | FWD  REV | AATGCTGAGCGACCACTCCATC  TCGGGTCATGTTCAAGTCCAGC |
| *FGF-1* | FWD  REV | CCAAGGAAACGTCCACAGTCAG  ACGGCTGAAGACATCCTGTCTC |
| *FGF-2* | FWD  REV | AAGCGGCTCTACTGCAAGAACG  CCTTGATAGACACAACTCCTCTC |
| *Angiopoietin-1* | FWD  REV | AACCGAGCCTACTCACAGTACG  GCATCCTTCGTGCTGAAATCGG |
| *Angiopoietin-2* | FWD  REV | AACTCGCTCCTTCAGAAGCAGC  TTCCGCACAGTCTCTGAAGGTG |

***Supplemental Figure 1.*** *(A) Representative ICC images of SSEA4 and OCT4 expression to confirm pluripotency of EV-producing iPSCs after 10 passages as well as absence of those markers from BDMSCs. (B) ICC images of SSEA4 and OCT4 expression to confirm pluripotency of iPSCs when cultured in “depleted” mTESR plus media. (C) Total particle count of samples subjected to the EV isolation process via TFF. mTESR plus media that hasn’t undergone “depletion” produces contaminants that are similarly sized to EVs. Upon depletion via ultracentrifugation (UC) for 16 hours at 100,000 x g, the depleted mTESR plus samples yield a particle count that is barely above the limit of detection (mTESR dep). This particle count is then increased upon culturing with iPSCs (iPSC dep). (D) Western blot analysis of iPSC EVs isolated via either TFF or UC demonstrate the presence of CD63 and absence of contaminating Calnexin. (D) Representative NTA distribution profiles of iPSC EVs isolated via UC or TFF.*

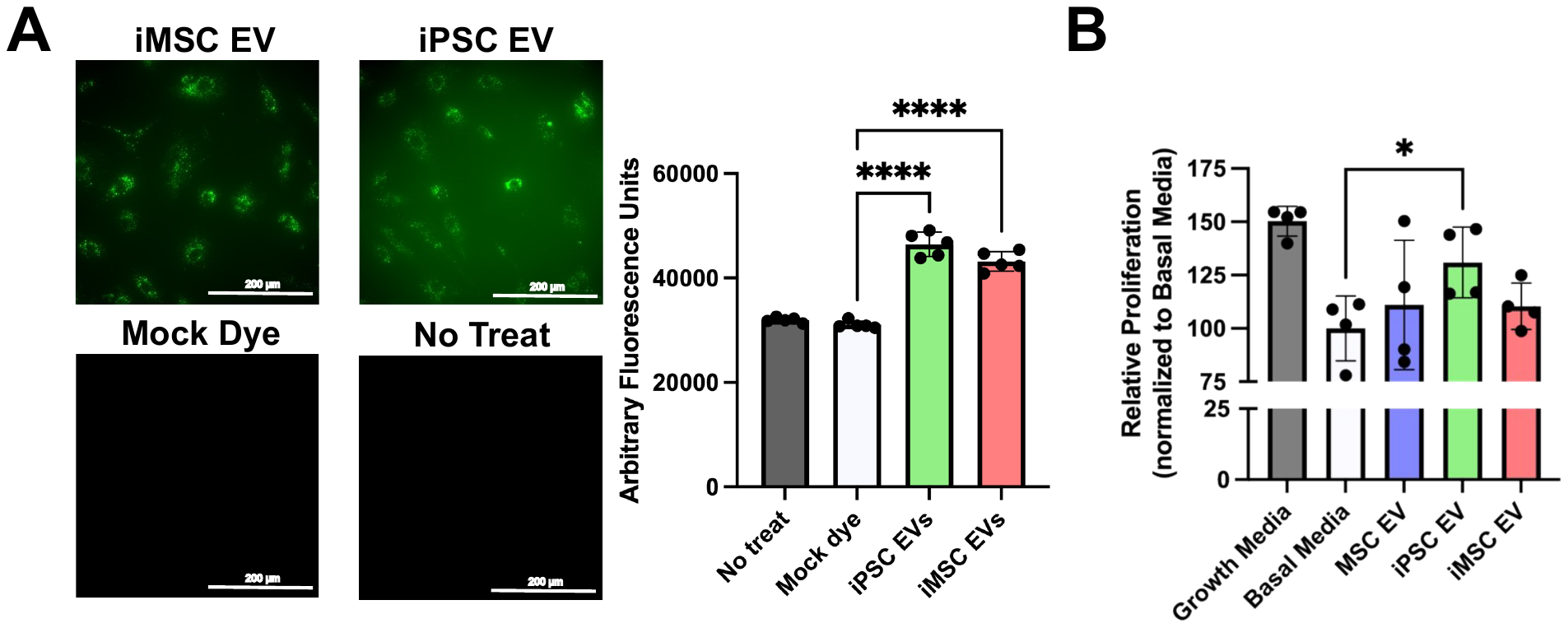

***Supplemental Figure 2.*** *(A) HUVECs were treated with PKH67 dyed EVs and imaged using a fluorescence microscope. Additionally, fluorescence intensity was quantified via plate reader (n=5). (B) Endothelial proliferation of HUVECs was quantified using a CCK8 assay after treatment with EVs in basal media (n=4). All values were expressed as mean ± standard deviation (*p < 0.05, ****p < 0.0001).*

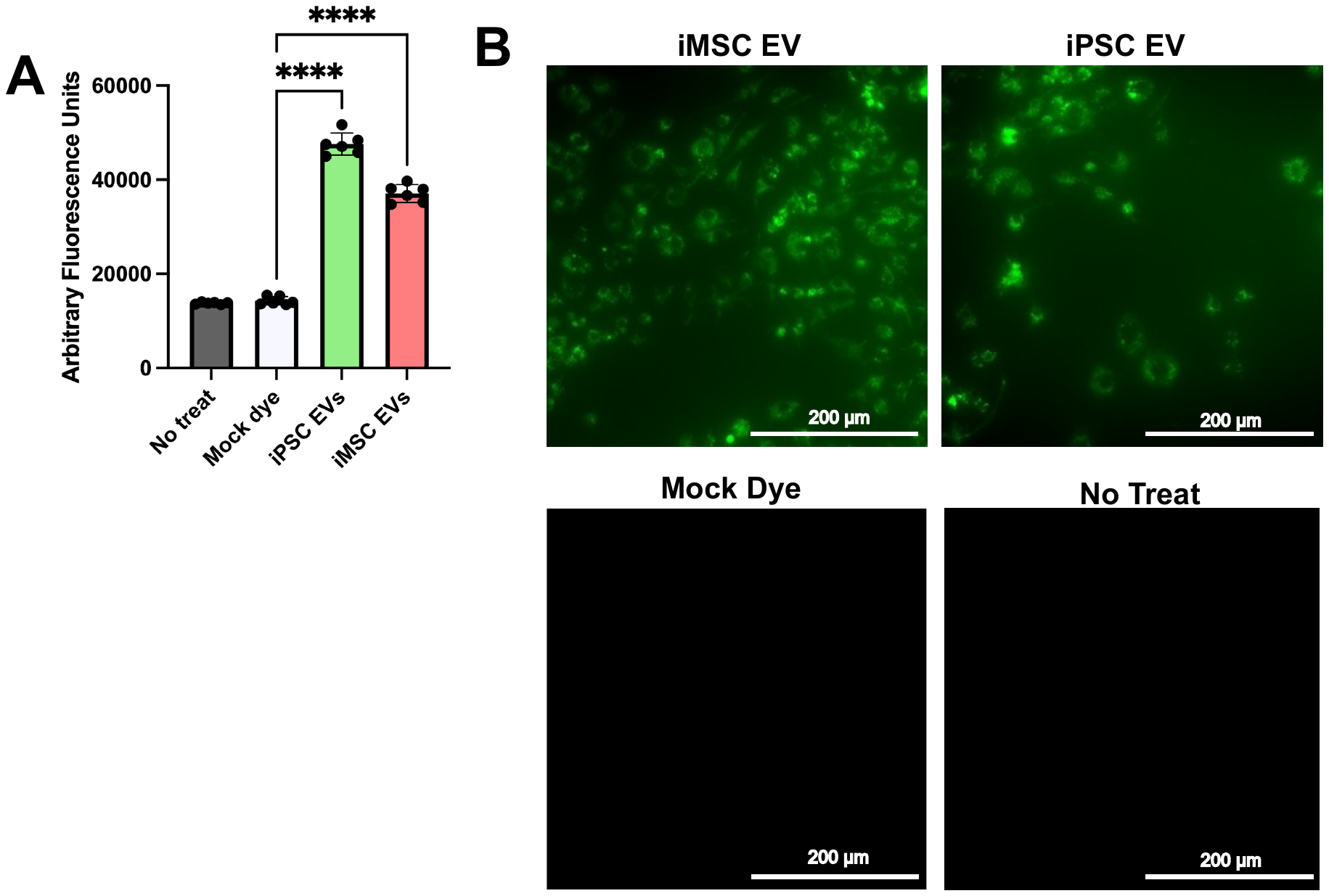

***Supplemental Figure 3.*** *RAW264.7s were treated with PKH67 dyed EVs and imaged with a fluorescence microscope. Fluorescence intensity was quantified using a plate reader (n=6). All values were expressed as mean ± standard deviation (****p < 0.0001).*

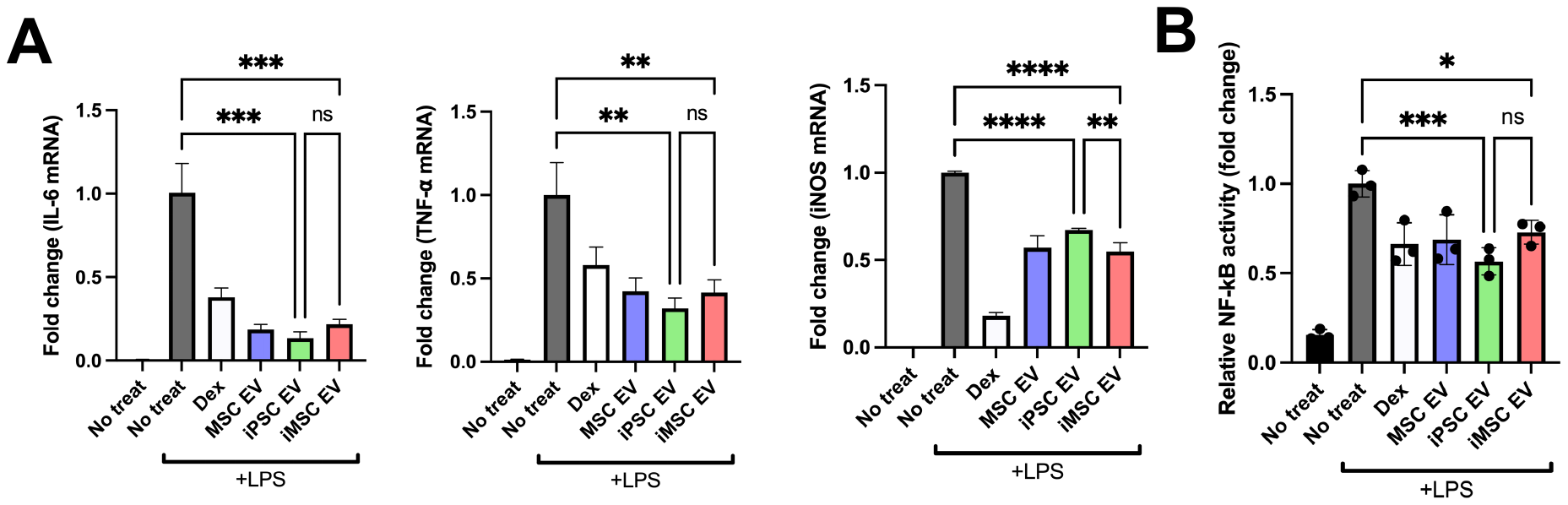

***Supplemental Figure 4****. (A) After EV treatment and LPS stimulation, RAW264.7 mouse macrophages were lysed and mRNA levels of pro-inflammatory cytokines/markers IL-6, TNF-⍺, and iNOS were quantified via RT-qPCR (n=3). (B) Using a RAW264.7 mouse macrophage Alkaline Phosphatase-based reporter cell line, NF𝛋B activity was quantified to determine inflammatory response on the transcriptional activator level (n=3). All values were expressed as mean ± standard deviation (*p < 0.05, **p < 0.01, ***p < 0.001, ****p < 0.0001).*

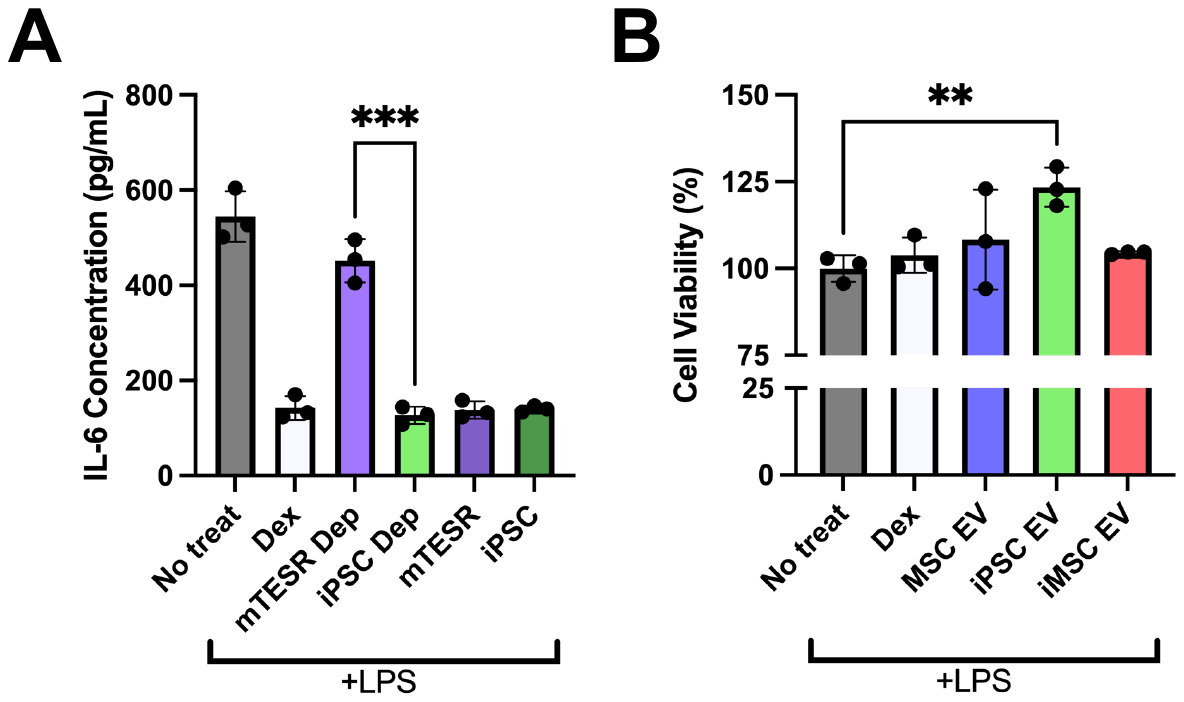

***Supplemental Figure 5.*** *(A) RAW264.7 mouse macrophages were pre-treated with samples that had undergone the EV isolation process at a dose of 5E9 particles/mL. The groups were 1. mTESR that had been depleted via UC for 16 hours (mTESR dep) 2. Conditioned media from iPSCs cultured with depleted mTESR media (iPSC dep) 3. Un-depleted mTESR media (mTESR) and 4. Conditioned media from iPSCs cultured with un-depleted mTESR media (iPSC). We observed that the resultant sample from un-depleted mTESR led to a substantial decrease in IL-6 secretion in our mouse macrophage assay, indicating that contaminants from the mTESR complete media, which are likely protein aggregates, induced an anti-inflammatory response. However, mTESR that had undergone a UC depletion process for large particle aggregates, did not lower IL-6 concentration. Additionally, upon culturing this depleted mTESR with iPSCs, the anti-inflammatory effect was rescued. This indicates that the EVs secreted by iPSCs indeed possess anti-inflammatory properties. (B) Again, using our RAW264.7 mouse macrophage assay with a pre-treatment regime, we looked to determine whether the decrease in inflammatory cytokine release was possibly due to a loss of macrophage viability/number upon treatment with iPSC EVs using a CCK8 assay. All values were expressed as mean ± standard deviation (**p < 0.01, ***p < 0.001).*

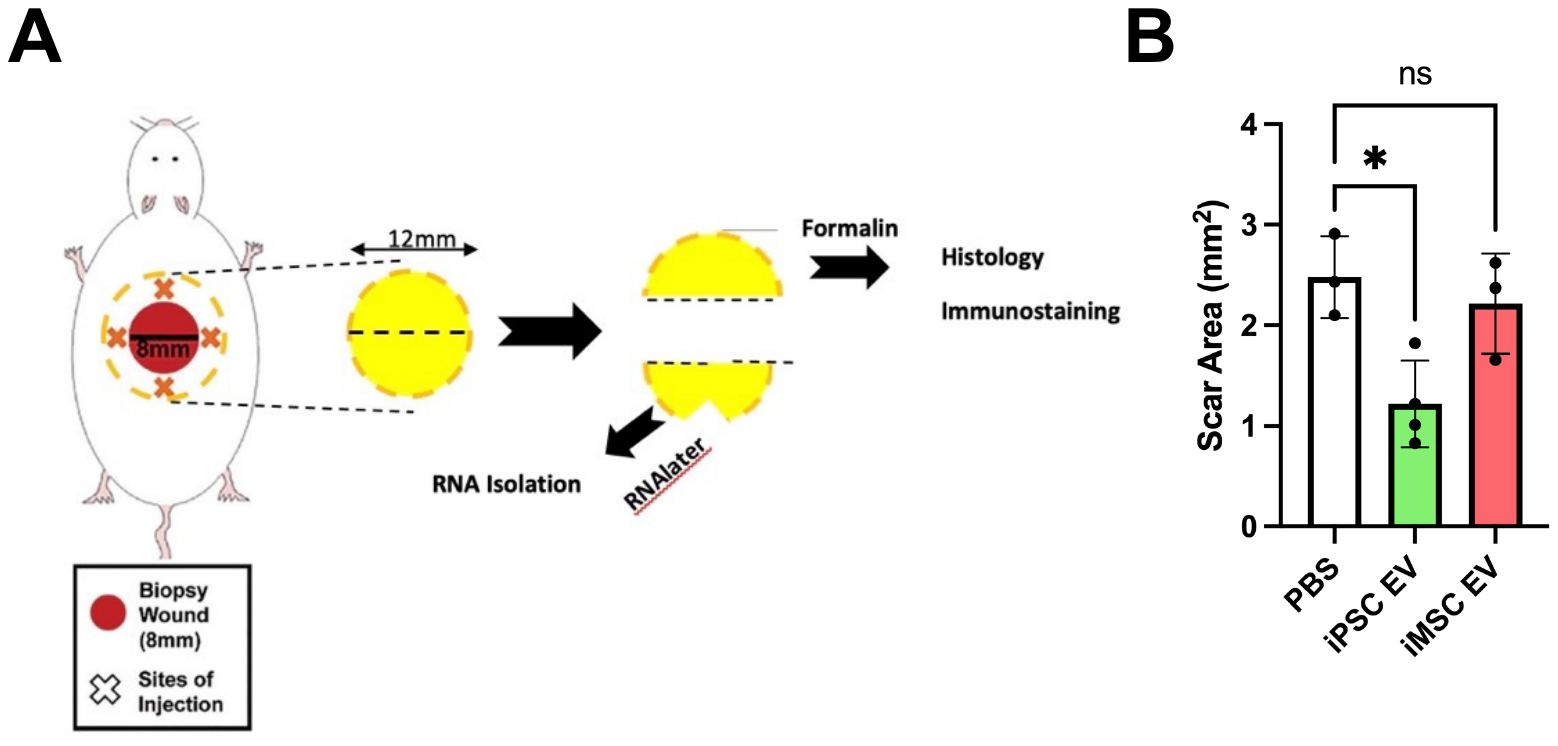

***Supplemental Figure 6.*** *(A) Schematic of wounding and EV injection procedure. An 8 mm punch biopsy was performed to create wounds and treatments were injected in a cross pattern surrounding the wound. At a later timepoint, a 12 mm punch biopsy was performed to extract tissue for both histological analyses as well as bulk RT-qPCR analyses. (B) Scar area was quantified from H&E-stained tissue slices from the wound bed by tracing the dermal granulation tissue (*p < 0.05).*

***Supplemental Figure 7.*** *(A)* *Representative IHC images of Ly6G stained tissues 3 days post EV injection. Ly6G fluorescence intensity over multiple fields of view was quantified and normalized to cell number obtained via DAPI staining. We see a non-statistically significant 30% decrease in Ly6G expression in wounds treated with iPSC EVs, indicative of lower neutrophil/monocyte/granulocyte infiltration. Surprisingly, we see a ~50% increase in infiltrate with iMSC EV treatment (n=4). (B) Inflammatory cytokine mRNA expression of iNOS and TGFβ was quantified via RT-qPCR from bulk mRNA in the wound bed 6 days post-wounding (n=4). (C) 18 days post-wounding, representative images of F4/80 IHC-stained tissue sections were taken, and F4/80 intensity was measured and normalized to cell number over multiple fields of view (n=4). (D) Representative images of CD206 IHC-stained tissue sections 18 days after initial wounding. Again, intensity was quantified and normalized to cell number (n=4). All values were expressed as mean ± standard deviation (*p < 0.05, **p < 0.01).*
